# The Nature of Genetic Susceptibility to Multiple Sclerosis

**DOI:** 10.1101/2020.08.13.249920

**Authors:** DS Goodin, P Khankhanian, PA Gourraud, N Vince

**Author notes:** Address for Correspondence: Douglas S. Goodin, MD, Department of Neurology, University of California, San Francisco, UCSF MS Center, 675 Nelson Rising Lane, Suite #221D, San Francisco, CA 94158, E mail. **Author Contributions** DSG: Conceptualized and led the project, analyzed and assisted in the interpretation of the data, developed the susceptibility-framework, and wrote the original draft of the manuscript. PK: Assisted in the critical interpretation of the data and review of the manuscript. PA: Assisted in the critical interpretation of the data and review of the manuscript. NV: Assisted in the analysis and critical interpretation of the data and the review of the manuscript.

## Abstract

**OBJECTIVE:** To explore the nature of MS-susceptibility and, by extension, other complex-genetic diseases.

**BACKGROUND:** Basic-epidemiological parameters of MS (e.g., prevalence, recurrence-risks for siblings and twins, time-dependent changes in sex-ratio, etc.) are well-established. Moreover, >200 genetic-loci are unequivocally MS-associated, especially the *HLA-DRB1*15:01~HLA-DQB1*06:02~a1* haplotype-association.

**DESIGN/METHODS:** We define the “genetically-susceptible” subset-(*G*) to include everyone with any non-zero life-time chance of developing MS. We analyze, mathematically, the implications that these epidemiological observations have regarding genetic susceptibility. In addition, we use the sex-ratio change (observed over a 35-year interval), to derive the relationship between MS-probability and an increasing likelihood of a suitable environmental-exposure.

**RESULTS:** We demonstrate that genetic-susceptibitly is restricted to less than 4.7% of populations across Europe and North America. Among carriers of the *HLA-DRB1*15:01~HLA-DQB1*06:02~a1* haplotype, fewer than 20% are even in the subset-(*G*). Women are less likely to be susceptible than men although their MS-penetrance is considerably greater. Response-curves for MS-probability increase with an increasing likelihood of a suitable environmental-exposure, especially among women. These environmental response-curves plateau at under 50% for women and at a significantly lower level for men.

**CONCLUSIONS:** MS is fundamentally a genetic disorder. Despite this, a suitable environmental-exposure is also critical for disease-pathogenesis. Genetic-susceptibility requires specific combinations of non-additive genetic risk-factors. For example, the *HLA-DRB1*15:01~HLA-DQB1*06:02~a1* haplotype, by itself, poses no MS-risk. Moreover, the fact that environmental-response-curves plateau below 50%, indicates that disease-pathogenesis is partly stochastic. By extension, other diseases for which monozygotic-twin recurrence-risks greatly exceed disease-prevalence (e.g., rheumatoid arthritis, diabetes, and celiac disease), must have a similar genetic basis.

**Author Summary:** We define a “genetic-susceptible” subset (*G*) of the general population (*Z*) to include everyone with any non-zero chance of developing MS over their life-time. Using well-established epidemiological data from across Europe and North America, we establish that genetic-susceptibility is confined to less than 4.7% of these populations. Thus, the large majority of individuals have no chance whatsoever of developing MS, irrespective of any environmental conditions that they may experience during their lifetimes. In this sense, MS is fundamentally a genetic disorder. And, indeed, more than 200 genetic-loci, in multiple genomic locations have now been well-established to be associated with MS. Notably, however, the *HLA-DRB1*15:01~HLA-DQB1*06:02~a1* or (*H+*) haplotype, which has, by far, the strongest MS-association of any, has a carrier frequency in the population of 23% in North America and Europe. Therefore, with genetic susceptibility in the population being less than 4.7%, more than 80% of (*H+*)-haplotype carriers, must not be genetically-susceptible and, thus, have no chance of developing MS. In this circumstance, genetic susceptibility to MS must arise from a combination of this haplotype with “susceptible states” at other genetic loci. By itself, the (*H+*)-haplotype poses no risk. Indeed, genetic-susceptibility, generally, seems to require specific combinations of non-additive genetic risk-factors.

Naturally, the conclusion that MS is fundamentally genetic does not preclude the possibility the environmental events are also critical to disease-pathogenesis. Using epidemiological data about the world-wide increase in the (*F:M*) sex-ratio for MS to construct (for men and women separately) the response curves relating an increasing likelihood of MS to an increasing likelihood of a sufficient environmental exposure (i.e., an exposure sufficient to cause MS in a susceptible individual). This analysis provides insight to both disease-susceptibility and disease-pathogenesis. First, men are more likely to be susceptible than women although susceptible women are considerably more likely to actually develop MS. Second, men seem to have a lower environmental threshold than women for developing MS. Nevertheless, women are more responsive to changes in the environmental conditions compared to men. Third, even with a maximal environmental exposure, susceptible women never exceed a 50% chance of developing MS. By contrast, susceptible men have a significantly lower likelihood (<10% chance) of developing MS. This indicates that stochastic factors must also be critical in disease pathogenesis.

Finally, the nature of genetic susceptibility developed here for MS is applicable to many other complex genetic disorders. Indeed, for any disease, in which the proband-wise *MZ*-twin concordance rate greatly exceeds the disease-prevalence in the population (e.g., type I diabetes, rheumatoid arthritis, and celiac disease), only a small fraction of the population can possibly be genetically susceptible, as defined.

## Introduction

The nature of susceptibility to multiple sclerosis (MS) is complex and involves both environmental and genetic factors [1–4]. Recently, considerable progress has been made in our understanding of the genetic basis for susceptibility to MS. Thus, to date, using genome-wide association screens (GWAS), which incorporate large arrays of single nucleotide polymorphisms (*SNPs*) scattered throughout the genome, over 200 common risk variants (located in diverse genomic regions) have been identified as being MS-associated [5–14]. Despite this recent explosion in the number of identified MS-associated regions, however, the association of susceptibility to MS with the *HLA-DRB1*15:01~HLA-DQB1*06:02* haplotype of the human leukocyte antigens (*HLA*) inside the major histocompatibility complex (*MHC*) has been known for decades [11,15–22]. We have recently identified an 11-*SNP* haplotype (*a1*), which adds further specificity to the description of this particular genetic association [23,24]. This *SNP*-haplotype spans 0.25 megabases (*mb*) of DNA surrounding the *HLA-DRB1* gene on the short arm of chromosome 6. It has the most significant association with MS of any *SNP*-haplotype in the genome, and it is tightly linked to the *HLA-DRB1*15:01~HLA-DQB1*06:02* haplotype [23,24]. For example, 99% of these (*a1*) *SNP*-haplotypes carry this *HLA*-haplotype and, conversely, 99% of these *HLA-*haplotypes carry the (*a1*) *SNP*-haplotype. In the Welcome Trust Case Control Consortium (WTCCC) dataset, the odds ratio (*OR*) for an association the full *HLA-DRB1*15:01~HLA-DQB1*06:02~a1* haplotype with MS was 3.28 (*p*≪10^−300^) and similar disease associations for portions of this haplotype have been consistently reported in many other MS populations across Europe and North America [11,15–22].

Despite the undoubted influence of genetic and environmental factors in MS-pathogenesis, susceptibility to MS might be envisioned in number of different ways. Four examples of disease states, for which we understand, generally, the pathophysiology, can be helpful to highlight some of the issues that might also be involved in MS pathogenesis.

First, sickle cell disease (*SCD*) occurs in ~3% of individuals in certain sub-Saharan regions of Africa [25]. All affected individuals are homozygous for the *HbS* mutation of the hemoglobin gene. Despite the fact that the clinical expression of *SCD* can be influenced by environmental factors such as strenuous exercise, high-altitude, infection, and dehydration, *SCD* is fundamentally a genetic disorder.

Second, each year, 5–20% of the population in North America gets the flu [25]. Although the genetic make-up might make one person more or less susceptible to a particular year’s variant, presumably, everyone could develop the flu if they had a sufficient exposure to the influenza virus. Therefore, despite the possible genetic differences in susceptibility, the flu is fundamentally an environmental (infectious) disease.

Third, the life-time probability of breast cancer in the US is ~12.5% in women and ~0.1% in men. Individuals (especially women) who carry the *BRCA1* or *BRCA2* mutations (<1% of the population) have 4-7 times the risk as that in the general population [25]. Nevertheless, presumably, there is a baseline risk of breast cancer such that no one is completely risk-free. Although the genetic make-up (including gender) influence the baseline risk and the environment likely affects the penetrance of the *BRCA* mutations, some breast cancer cases are fundamentally genetic and others are fundamentally environmental (of unclear type, but possibly due to exposures such as by toxins, radiation, pregnancy, or other occurrences).

Fourth, the human immunodeficiency virus (*HIV*) can infect anyone in the population although individuals who engage in certain high-risk behaviors (e.g., having unprotected anal-receptive sex or using IV drugs and sharing needles) are particularly susceptible [25]. Among persons of northern European extraction, ~1% are homozygous for the *Δ-32 mutation* of the *CCR5* gene and are almost completely resistant to *HIV*. Consequently, *HIV* infection is fundamentally an environmental disorder (infectious) with an interaction between two environmental factors (i.e., the virus and specific high-risk behaviors). However, certain genetic traits (*e.g., the Δ-32 mutation*) can be decisive in determining the degree of susceptibility.

Whether susceptibility to MS resembles any of these disease-states (or some other) is unknown although its polygenic nature is certain [5–14]. Nevertheless, several recent epidemiological observations in MS bear directly on the different possibilities. In this paper, we utilize directly observable, and well-established, “population parameters” (e.g., twin concordance rates, the percent women among MS patients, the population prevalence of MS, time-dependent changes in the gender-ratio, etc.) to logically infer the values of other non-observable parameters of interest (e.g., the population probability of being genetically susceptible, the percentage of susceptible individuals who are women, the likelihood that a susceptible individual receives a sufficient environmental exposure, etc.).

## Methods

For the purpose of this analysis we define, explicitly, five general terms (*Table 1*). The first term is {*P(MS)*}, which represents the expected life-time probability that a random individual from the general population (*Z*) will develop MS {i.e., the expected penetrance: *P*(*MS*) = *P*(*MS* | *Z*)}. As discussed below, this parameter is related to, but not the same as, the population prevalence.

**Table 1.** Definitions for Epidemiological Parameters used in the Analysis*

| Parameter | Definition |
| --- | --- |
| $(Z)$ | Set of all individuals in the population |
| $P(MS)$ | Expected life-time probability of developing *MS* for a member of $(Z)$ |
| $(G)$ | Subset of all individuals in $(Z)$ who have any non-zero chance of getting *MS* |
| $(G-)$ | Subset of all individuals in $(Z)$ who have no chance of getting *MS* |
| $(G1)$ | For a bimodal distribution, a subset of “high-penetrance” individuals in $(G)$ |
| $(G2)$ | The subset of “low-penetrance” individuals in $(G)$ such that: $(G1) + (G2) = (G)$ |
| $\{X\}$ | Set of all penetrance values for members of the subset $(G)$ |
| $(H+)$ | Set of all carriers of the *DRB1\*15:01~DQB1\*06:02~a1* haplotype in $(Z)$ |
| $(H-)$ | Set of all non-carriers of the $(H+)$ HLA haplotype in $(Z)$ |
| $(M)$ | Set of all men in $(Z)$ |
| $(F)$ | Set of all women in $(Z)$ |
| $P(E)$ | Expected probability of an environmental exposure “sufficient to cause *MS*” in the set $(G)$, given the prevailing environmental conditions of the time |
| $(E_1)$ | That part of the sufficient environmental exposure shared exclusively by *MZ*- or *DZ*-twins – especially during the *IU* and early post-natal period |
| $(E_2)$ | That part of the sufficient environmental exposure shared by the population generally: |
| $(E_3)$ | The potential part of a sufficient environmental exposure due exclusively to the shared microenvironment of families. However, observationally [62-68]:<br>$P(E) = P(E_1, E_2, E_3) = P(E_1, E_2)$ |
| $P(MS \vert MZ_{MS})$ | Expected life-time probability of developing *MS* for an *MZ*-twin whose co-twin either has, or will develop, *MS* |
| $P(MS \vert DZ_{MS})$ | Expected life-time probability of developing *MS* for a *DZ*-twin whose co-twin either has, or will develop, *MS* |
| $P(MS \vert S_{MS})$ | Expected life-time probability of developing *MS* for a sibling whose co-sibling either has, or will develop, *MS* |
| $P(MS \vert IG_{MS})$ | Value of $P(MS \vert MZ_{MS})$, which has been adjusted to exclude the impact of the similar *IU* and early post-natal environments of *MZ*-twins |
| $P(MZ_{MS})$<br>$P(IG_{MS})$ | Expected life-time probability of developing *MS* for an individual from any *MZ*-twin-ship. $P(MZ_{MS}) = P(IG_{MS}) = P(MS)$ |

The second term is {*P*(*G*)}, which represents the expected probability that a random individual from (*Z*) is also a member of the subset-(*G*) – i.e., *P*(*G*) = *P*(*G* | *Z*). In turn, we define the subset-(*G*) to include everyone who has any non-zero chance of developing MS (i.e., regardless of how small that risk might be). We also define the set {*X*} to be the set of penetrance values for members of the subset-(*G*). In our analysis, we will only consider the possibility that set {*X*} has either a unimodal or a bimodal distribution. We will discount the possibility that {*X*} has a more extreme distribution such as one that is trimodal or multimodal in nature.

The third term {*P*(*E*)}, represents the expected probability that a member of the (*G*)-subset will experience an environmental exposure sufficient to cause MS given the prevailing environmental conditions of the time.

The fourth is a set of related terms {*P*(*MS*│*MZ*_*MS*_), *P*(*MS*│*DZ*_*MS*_), and *P*(*MS*│*S*_*MS*_)}. The 1^st^ two terms {*P*(*MS*│*MZ*_*MS*_)} and {*P*(*MS*│*DZ*_*MS*_)} represent the expected conditional life-time probability of developing MS for an individual from either a monozygotic (*MZ*) or a dizygotic (*DZ*) twin-ship, given the fact that their co-twin either has or will develop MS. These probabilities are estimated by the observed proband-wise concordance rate for either *MZ*-twins or *DZ*-twins [26]. In a similar manner, the term {*P*(*MS*│*S*_*MS*_)} represents the expected conditional life-time probability of developing MS in a sibling (*S*), given the fact that their co-sibling either has or will develop MS.

Lastly, the term {*P*(*MS*│*IG*_*MS*_)} represents the adjusted proband-wise concordance rate for *MZ*-twins. Such an adjustment may be necessary because concordant *MZ*-twins, in addition to sharing their identical genotypes (*IG*), also share similar intrauterine (*IU*) and early post-natal environments. Thus, it is possible that these shared early environmental experiences of twins might significantly impact the likelihood of their developing MS in the future. One method to estimate the adjustment necessary in such a circumstance is to consider the difference in concordance rates between non-twin siblings and fraternal twins (i.e., siblings who share the same genetic relationship but who are divergent in their *IU* and early post-natal experiences). Although epidemiological studies have differed somewhat with regard to the magnitude of any such differences [27–34], population-based studies out of Canada suggest that the impact of these early environmental events may be substantial [29]. As demonstrated in the *Supplemental Material*, we can use the observed recurrence-rate data (*Table 2*) to make this adjustment such that:

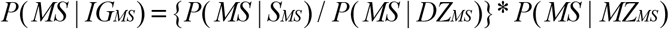

**Table 2.** Canadian Population Data on *MZ*-Twin Concordance broken down by (*H+*)-haplotype and Gender-Status *

| ***HLA-DRB1\*15 Status*** | ***H+*** | ***H-*** | ***Totals*** |
| --- | --- | --- | --- |
| Concordant for MS (C) | 9 | 11 | 20 |
| Discordant for MS (D) | 31 | 42 | 73 |
| Totals | 40 | 53 | 93 |
| Pair-wise Concordance | 0.23 | 0.21 | 0.22 |
| Proband-wise Concordance | 0.31 | 0.29 | 0.30 |
| ***Gender Status*** | ***Women*** | ***Men*** | ***Totals*** |
| Concordant for MS (C) | 22 | 2 | 24 |
| Discordant for MS (D) | 66 | 43 | 109 |
| Totals | 88 | 45 | 133 |
| Pair-wise Concordance | 0.25 | 0.04 | 0.18 |
| Proband-wise Concordance | 0.34 | 0.07 | 0.25 |

***Summary of Canadian Data:***
\* Data from Willer et al. [29] and Orton et al. [54]
– the MZ-twins were drawn from the 19,938 MS-patients in the CCGPSMS database
Pair-wise concordance calculated as: $C/(C+D)$
Proband-wise concordance calculated as: $2C/(2C+D)$
– adjusted [26] for double ascertainment (13/24=54%)
\*\* D Sadovnick (personal communication). Based on 400 Controls. Rate confirmed in a large transplant database.

From these definitions and relationships, we can use well-established values for the different population parameters to logically deduce the value of the another, non-observable, parameter {*P*(*MS* | *G*)}, which represents the conditional life-time probability of getting MS for a member of the (*G*)-subset. This term is referred to as the expected penetrance for the (*G*)-subset.

We note that, from the definition of the (*G*)-subset, everyone who actually develops MS during their life-time must belong to this subset. Therefore, the joint probability {*P*(*MS, G*)} must be the same as {*P(MS)*}, so that, by definition:

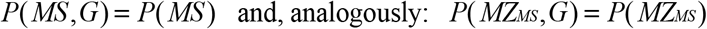

From this, and from the definition of conditional probability:

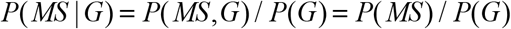

This equation can be re-arranged to yield:

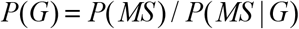

This relationship, once established, can then be used to assess the fundamental nature of MS pathogenesis. For example, if {*P*(*G*) = 1}, then anyone can get the disease under the right environmental circumstances (*e.g., flu, breast cancer, & HIV*) and we would conclude that MS must, in some cases, be caused by purely environmental factors. Notably, however, such circumstance does not preclude the possibility that genetic factors strongly influence the likelihood of disease (*e.g., breast cancer & HIV*).

By contrast, if {*P*(*G*) < 1}, this indicates that only certain individuals can possibly get the disease (*e.g., SCD*) and that, therefore, MS must be fundamentally a genetic disorder (i.e., unless a person has the correct genetic make-up, they have no chance, whatsoever, of getting the disease, regardless of their environmental exposure). Naturally, also, such a conclusion would have no bearing on whether disease pathogenesis also requires the co-occurrence of specific environmental events. Also, in this circumstance, how we might characterize the nature of genetic susceptibility, would depend upon the degree to which *P*(*G*) was less the unity and upon the magnitude of the disparity between any so-called “high” and “low” penetrance subgroups. For example, in *HIV*, if homozygous *Δ-32* mutations were completely protective, then: *P*(*G*) = 0.99. In this circumstance, however, we would likely characterize the disease as being fundamentally environmental and the homozygous *Δ-32* mutations as being protective rather characterizing every other genotype as being “susceptible”. By contrast, in *SCD*, where: *P*(*G*) = 0.03, we would characterize carrying homozygous *HbS* mutations as the defining trait for membership in the “genetically susceptible” subset. Even if it were possible, in extremely rare circumstances, for an individual to develop *SCD* in the absence of homozygous *HbS* mutations, we would still consider this disease to be fundamentally genetic.

## Results

### 1. MS Penetrance in the General Population – *P*(*MS*)

***Conclusion:*** *P*(*MS*) ≈ 0.003

***Argument:*** One possible estimate of *P(MS)* could be the prevalence of MS in a population. However, because the clinical onset of MS occurs largely between the ages of 15 and 45 years (e.g., *Fig 1*), the measured cross-sectional prevalence of MS (using the entire population as the denominator) will necessarily include individuals with different likelihoods of having already developed MS [36]. For example, using the 2010 United States census data (for the total resident population – see *Fig 2*) as an approximation, we can divide the general population (*Z*) into the three mutually exclusive age-bands (*A1, A2,* and *A3*), such that:

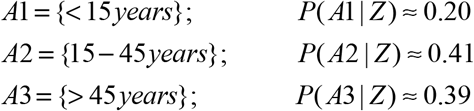

**Figure 1.**
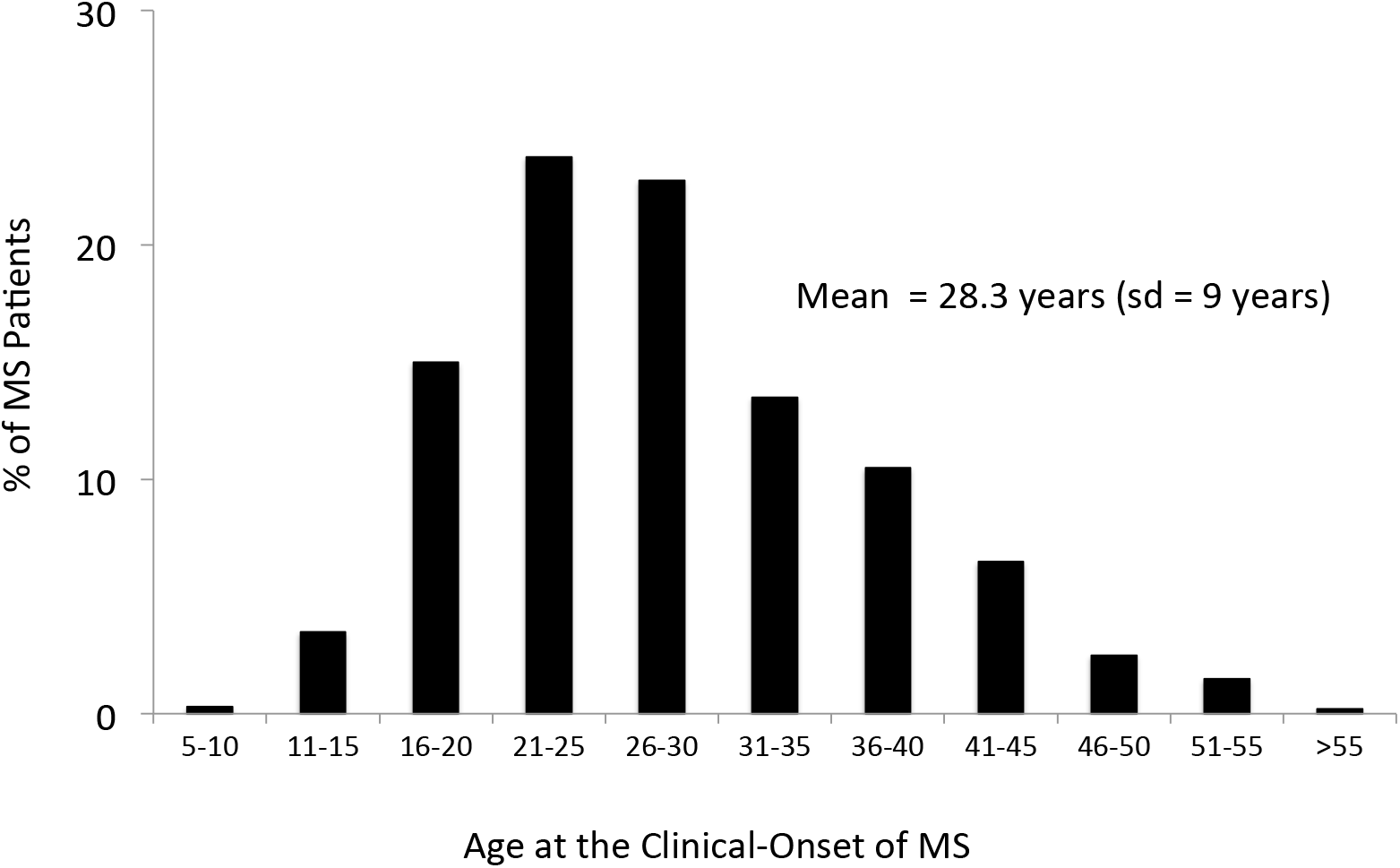
The distribution of the age at the clinical-onset of disease in a cohort of 1,463 patients with MS (sd=standard deviation). *Data from Liguori et al. [36*].

**Figure 2.**
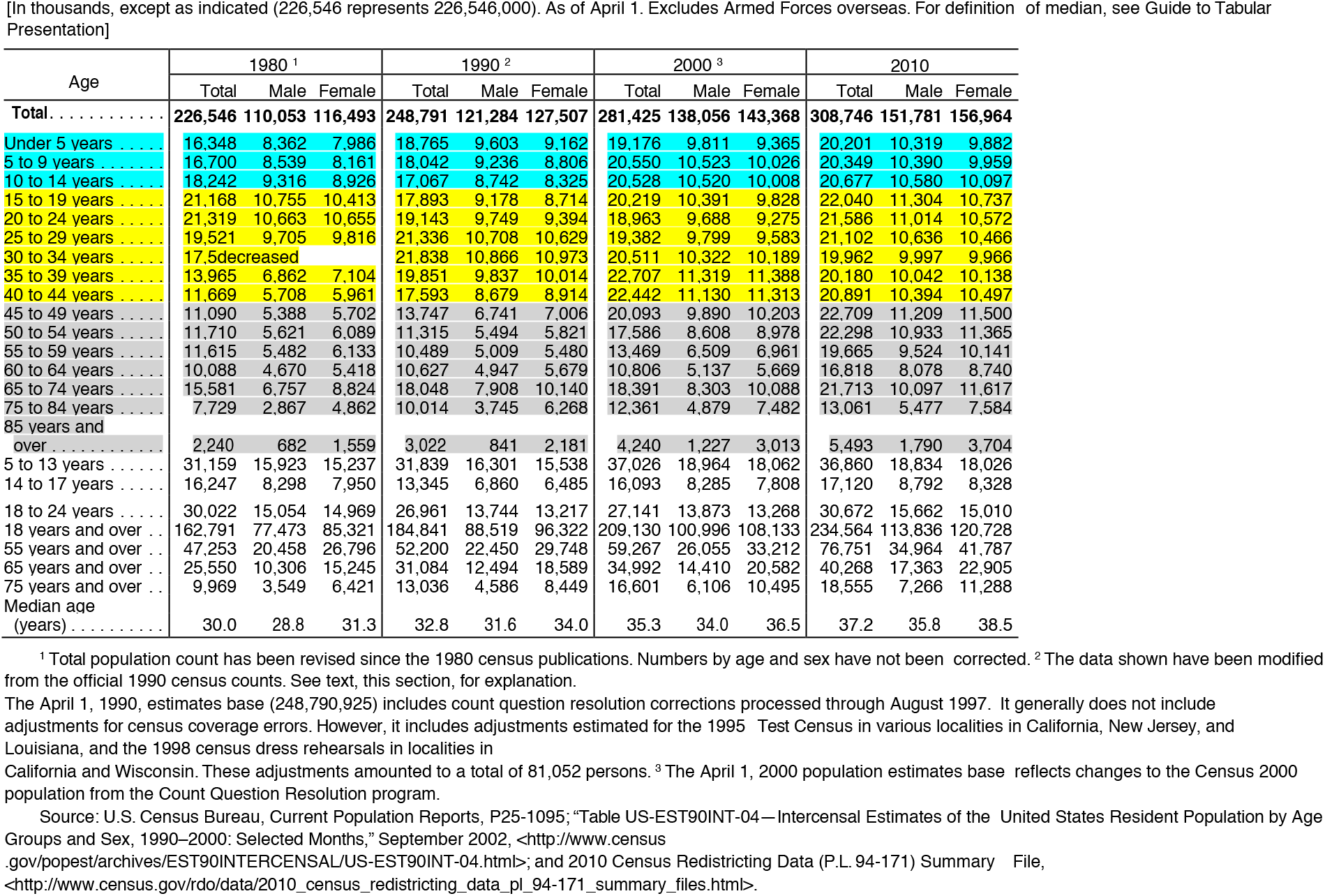
US Census Data (for each decade from 1980 to 2010) for Resident Population by Age and Sex. Age-band (*A1*) is highlighted in turquoise; Age-band (*A2*) is highlighted in yellow; and Age-band (*A3*) is highlighted in grey.

Because so few of MS patients have their disease onset prior to the age of 15 years (e.g., *Fig 1*) it seems a reasonable approximation that:

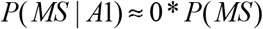

By contrast, as noted above, the age group (15–45 years) accounts for the large majority of clinical onsets, which have a roughly symmetrical distribution with a mean of 28.3 years (*Fig 1*). If the distribution were exactly symmetrical and centered on 30 years, the measured prevalence in this age band would be ~50% of the value of *P*(*MS*). Therefore, it seems reasonable to estimate:

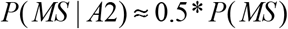

For the older age band (>45 years) most patients will have already developed the disease (*Fig 1*). Thus, on the one hand, one might expect that the measured prevalence in this age-band to be equal to *P(MS)*. On the other hand, there is a small but definite excessive mortality in MS such that life expectancy is reduced in MS-patients by about 5–10 years [37–41]. This will make the estimate too small by some amount. However, it seems unlikely for this reduction to be more than 25%. Thus, a range of plausible estimates is likely to be:

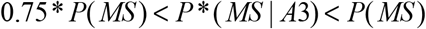

Combining these three different estimates yields the estimate:

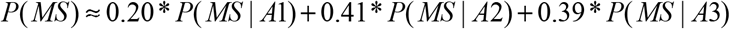

Defining the measured prevalence in the population as (*prev*), this estimate translates to:

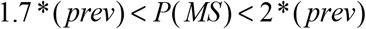

A second method to estimate *P*(*MS*) would be to use a measured prevalence for MS, which is restricted to the age-band of 45–54 years. Thus, within this age-band, almost all patients will have already experienced their clinical-onset and only a few will have experienced their (expected) excessive mortality. Consequently, by this method:

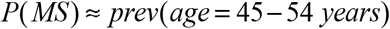

A third method would be to use population-based death data and to consider the percentage of death certificates that mention the diagnosis of MS (not necessarily as, but including, the immediate, underlying, or contributing cause). Thus, by the time of death, any case of clinically evident MS must, by definition, have already declared itself. Consequently, by this method:

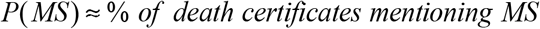

In 2001, we took a cursory (*unpublished*) look at the Kaiser northern California database. At the time, there were 4,352 unique persons in the database with a diagnostic code for MS. With 2.9 million persons enrolled in Kaiser northern California at the time, and if this population is a representative sample, this would translate to an MS-prevalence in northern California of 150 per 100,000 population. Such an estimate is consistent with many other published studies in northern populations, which generally find the prevalence of MS to be 100–200 per 100,000 population [42].

Similarly, in a Swedish study by Sundström and co-workers [43], the age-specific prevalence of MS in the 45-54 year age-band was reported to be 304 per 100,000 population.

And, finally, in a recent population-based multiple-cause-death study from British Columbia [44], a diagnosis of MS was mentioned on 0.28% of the death certificates.

Thus, all three of these methods of estimation are quite consistent with each other and support the conclusion that, in the northern populations of Europe and the Americas:

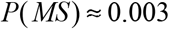

However, despite the notable consistency of these three estimates, each of these methods relates only to “diagnosed” MS in the general population (*Z*). However, if undiagnosed (i.e., pathological) MS is included in the calculation [45–48], this estimate may increase by as much as 50-100% (*see #8 below*).

### 2. Adjusting for the Shared Early Environment of *MZ-*twins – *P(MS*│*IG_MS_)*

***Conclusion:*** *P*(*MS* | *IG*_*MS*_) = *P*(*MS | G*, *IG*_*MS*_) ≈ 0.134

***Argument:*** Most epidemiological studies in northern populations report the proband-wise concordance rate for *MZ-*twins to be in the range of 25–30% [27–35]. Using the population data out of Canada (*Table 2*), leads to the estimate of:

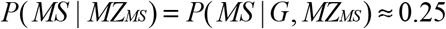

Suppose that each of the (*k* = 1,2,…, *n*) individuals within the general population (*Z*), has a unique genotype (*G*_*k*_) such that:

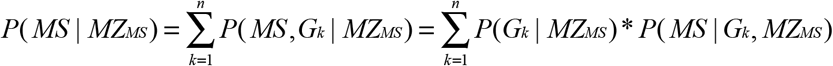

We can then define the term (*IG*_*MS*_) such that:

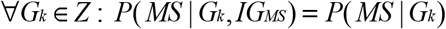

where:

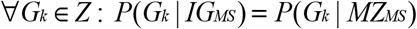

Therefore:

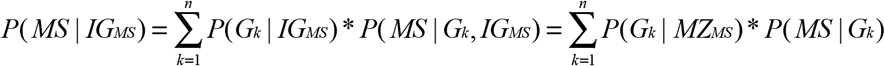

This is just the expected “adjusted” penetrance for the (*MZ*_*MS*_) subset. As discussed earlier and, as developed in the *Supplemental Material*, {*P*(*MS* | *IG*_*MS*_)} can be estimated from the difference in proband-wise concordance rates between siblings and fraternal twins. Using the Canadian population-based data (*Table 2*) on the recurrence risks in non-twin siblings and *DZ*-twins (concordance rates for siblings=2.9%; concordance rates for *DZ*-twins=5.4%) to make this adjustment (*see above*) leads to the estimate of:

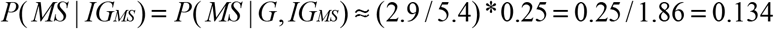

### 3. Proposition #1

a. If the individual values for penetrance within the (*G*)-subset are distributed in a unimodal manner, then:

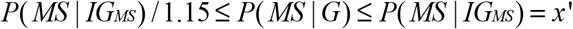
b. If these penetrance values are distributed in a bimodal manner, and if: *P*(*MS* | *G*) ≥ *x*′/ 2, then the Upper Solution limits always apply.
c. And finally, for more extreme bimodal distributions – i.e., *P*(*MS* | *G*) < *x*′/ 2 – then the Lower Solution applies.

***Proof:*** Among the (*n*) individuals in the general population (*Z*), we have already defined the subset-(*G*), which consists of everyone who has any non-zero life-time probability of developing MS. Thus, each of the (*m*) individuals in the (*G*)-subset (*i* = 1,2,…, *m*) has a unique genotype (*G*_*i*_), such that:

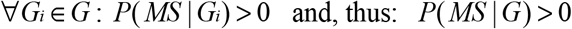

For ease of notation, we define two variables, (*x*_*i*_) and (*x*), such that:

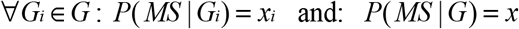

Thus, (*x*_*i*_) represents the expected penetrance for MS in the *i*^*th*^ individual of the (*G*)-subset. Even if this penetrance exactly matches that of another person, (*x*_*i*_) is still unique to the *i*^*th*^ individual. Also, considering each of the members of the (*G*)-subset, we can define the set {*X*} such that:

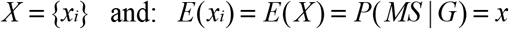

Because the subset-(*G*) forms a partition of the population (*Z*), each of the (*m*2 = *n* − *m*) individuals, who are not in the (*G*)-subset, belongs to the mutually exclusive “non-susceptible” subset (*G*−). Moreover, each of the (*j* = 1,2,…, *m*2) individuals in the (*G*−)-subset has a unique genotype (*G*_*j*_), which has a zero conditional life-time probability of developing MS, so that:

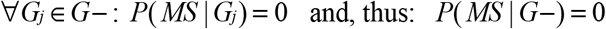

Also, we can partition the subset (*G*) into two mutually exclusive sub-subsets, (*G*1) and (*G*2) such that the sub-subset (*G*1) has a penetrance as great or greater than that of the sub-subset (*G*2). Again, for ease of notation, we further define the quantities (*x*′, *x*_1_, *x*_1_′, *x*_2_, & *x*_2_′) such that:

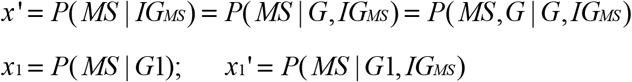

and:

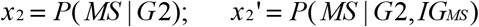

In earlier iterations of this analysis [3,4,49,50], we defined the subset-(*G*) differently – i.e., ∀*G*_*i*_ ∈*G* : *P*(*MS* | *G*_*i*_) ≥ *P*(*MS*). We have chosen the current definition because it considerably simplifies the biological interpretation of the findings. Nevertheless, we note that, when {*P*(*G*) = 1}, the new (*G*1)-subset is, effectively, identical to the (*G*)-subset defined earlier (*see below*).

We define the term {*P*(*MZ*_*MS*_)} to represent the life-time probability of developing MS for any single individual from an *MZ* twin-ship (i.e., where the status of their co-twin is unknown). Because identical twinning is considered non-hereditary [51], we expect that:

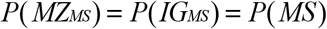

As noted earlier, we also define the set {*X*} set to consist of the individual *MS*-penetrance values for all members of the (*G*)-subset. Thus, the variance 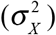 of the set {*X*} can be expressed as:

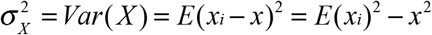

Also:

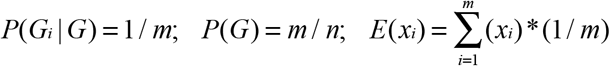

and:

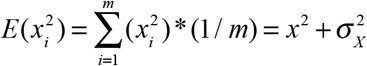

It follows directly from the definitions of {*P*(*G*)} and {*P*(*MS*│*IG*_*MS*_)} – *see #2 above* – that:

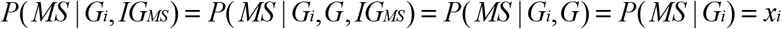

1. Therefore, the probability {*P*(*MS*,*G*_*i*_ | *G*, *IG*_*MS*_)} can be re-written as:

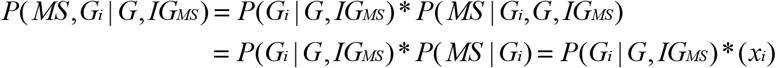
2. In turn, the term *P*(*G*_*i*_ | *G*, *IG*_*MS*_) can be re-written as:

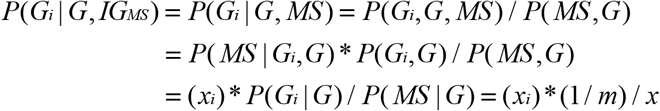

Combining these two Equations (*i.e., 1 & 2 above*) yields:

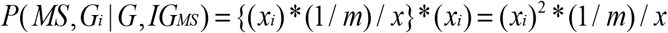

However:

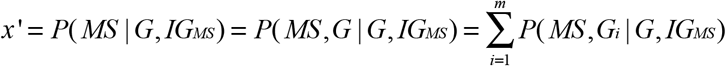

Where:

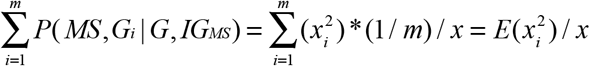

Consequently:

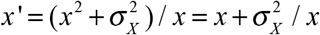

and, with rearrangement:

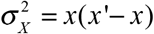

Notably, this equation can also be rearranged to yield a quadratic in (*x*) of:

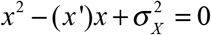

In turn, this quadratic equation can be solved to yield:

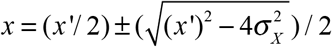

which has real, non-negative, solutions only for:

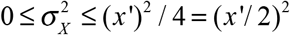

The maximum variance for <u>any</u> distribution [52,53] on the closed interval [*a*,*b*] is:

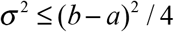

Consequently, the maximum variance from the quadtatic is the same that for the interval [0, *x*′], which is:

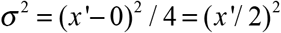

In addition, this maximum variance, (*x*′/ 2)^2^, occurs when the distribution of individual penetrance values in the set {*X*} is bimodal [52,53], such that half of the (*G*)-subset has a penetrance of (0) and the other half has a penetrance of (*x*′). At this point, for each of the two quadratic solutions:

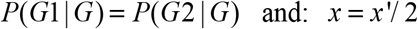

From this point the variance of the (*G*)-subset decreases both when:

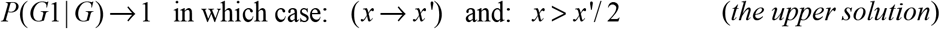

and when:

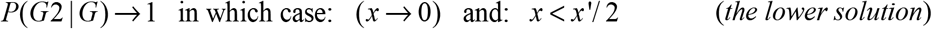

By definition, any solution requiring {*P*(*MS* | *G*_*i*_) = 0} for any portion of (*G*) is excluded.

Therefore, the upper solution becomes:

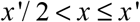

And the lower solution becomes:

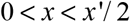

Moreover: when: 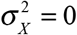; then: *x* = *x*′

#### The Upper Solution

The upper solution, as: (*x* → *x*′), represents the gradual transition from a bimodal distribution to a unimodal distribution and, ultimately, to a distribution, in which every genotype in (*G*) has exactly the same penetrance (i.e., *x* = *x*′). As noted earlier (*above*), the upper solution requires that:

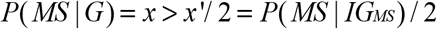

Also, as demonstrated by others [52], the maximum variance of <u>any</u> unimodal distribution on the closed interval [*a*,*b*] is: σ^2^ ≤ (*b* − *a*)^2^ / 9. Considering a unimodal distribution in the interval (0, *x*′], therefore:

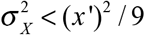

Substituting this limit into the upper quadratic solution (*above*) – assuming this limit applies equally to the set {*X*} – yields:

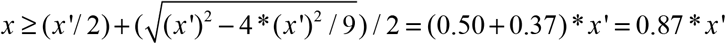

Consequently, for a unimodal distribution:

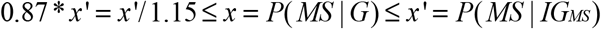

#### The Lower Solution

By contrast, the lower solution as: (*x* → 0), represents an increasingly assymetric bimodal distribution of penetrance values within the (*G*)-subset where: {*P*(*G*2 | *G*) > *P*(*G*1| *G*)}. As noted above, for this lower solution to apply, requires that:

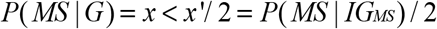

Notably, the value of (*x*′) represents an observed population parameter and is, therefore, “fixed”, allowing for the possiblity of error in its observation. Even with this constraint, however, some Lower Solution distributions are possible, which meet both of these two requirements and, for which, *P*(*G*) = 1 (*see #4, below*)

#### Breast Cancer

As an example, it is instructive to apply this same analysis to the risk in women of developing breast cancer (*descsribed briefly in the Introduction*). Clearly, this distribution is bimodal with <1% of women possessing the *BRCA* mutations, and with these individuals having 4–7 times the risk of breast cancer as that for everyone else. For this analysis, we assume that the subsets of women with (*G*1) and without (*G*2) *BRCA* mutations have a uniform penetrance within each subset. Also, we will also use parameter values that conform to the known epidemiology of breast cancer in women (*BC*) such that:

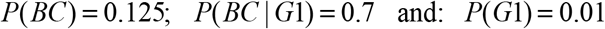

Under these conditions, and in all circumstances, it is the case that:

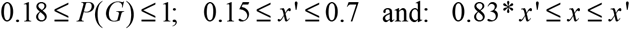

Although, unlike MS, we don’t have “observational” estimates for adjusted the *MZ*-twin recurrence risk (*x’*), these circumstances for breast cancer, clearly, conform to the upper solution of the quadratic equation (*above*). For example, if this recurrence risk were (~15%) then:{*P*(*G*) = 1} and: {*x* = 0.83* *x*′}. In this case, the fact that the distribution is bimodal is confirmed by the fact that the value of (*x*) is below the lower limit for a unimodal distribution (*see above*). By contrast, if all breast cancers are, to some degree, purely genetic disorders – {i.e., if: (*P*(*G*) < 1)} – then, as *P*(*G*) decreases, the value of (*x*) will increase. Nevertheless, the bimodality of the distribution will still be evident down to *P*(*G*) = 0.86. Below this point, however, the bimodal nature of the distribution will no longer be distinguishible (purely by consideration of the variance) from a unimodal distribution. Regardless, however, using these parameter values, the distribution would not actually become unimodal until the point at which: {*x* = *x*′}.

### 4. Genetic Susceptibility to MS (not considering gender)

***Conclusions:*** 0.02 ≤ *P*(*G*) ≤ 0.045

***Argument:*** From the Upper Solution in *Proposition #1* and in conjunction with our estimate for {*P*(*MS* | *IG*_*MS*_)}, it follows directly, that:

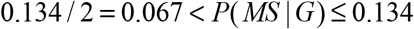

Using the relationship developed in the *Methods* of:

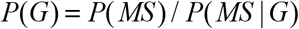

With this we have all the data necessary to establish the limits for the percentage of the population who are members of the (*G*)-subset. Thus, using this range for *P*(*MS*│*G*), together with our estimate for *P*(*MS*) – *see #1 above* – it follows that:

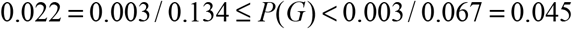

Consequently, by this analysis, only 4.5% or less of the general population (*Z*) could possibly be genetically susceptible to getting MS and the remainder of the population would have no possibility of getting this condition, regardless of their environmental experiences. Multiple reports from other MS-populations throughout Europe and North America yield very similar Upper Solution estimates for *P*(*G*), which seems to be independent of latitude (*Table 3*).

**Table 3.** MS prevalence, *MZ*-twin concordance {*P*(*MS│MZ_MS_)*}, and “genetic susceptibility” {*P*(*G*)} – for the Upper Solution (*see Text*) – in different geographical locations.

| Geographical Location | MS Prevalence<sup>*</sup> | $P(MS\vert MZ_{MS})^{\dagger}$ | Latitude | $P(G)^{\dagger\dagger}$ |
| --- | --- | --- | --- | --- |
| ***North America*** |  |  |  |  |
| Canada [29] | 68 – 248 | 0.253 | N45-60° | 0.01 – 0.07 |
| Northern US [32] | 100 – 160 | 0.314 | N41-45° | 0.01 – 0.04 |
| Southern US [32] | 22 – 112 | 0.174 | N30-41° | 0.005 – 0.05 |
| ***Europe*** |  |  |  |  |
| Finland [34] | 52 – 93 | 0.462 | N60-70° | 0.004 – 0.015 |
| Denmark [30,31] | 110 | 0.240 | N55-58° | 0.017 – 0.03 |
| British Isles [28] | 74 – 193 | 0.400 | N50-59° | 0.007 – 0.04 |
| France [27] | 32 – 65 | 0.111 | N44-50° | 0.01 – 0.04 |
| Sardinia [33] | 144 – 152 | 0.222 | N39-41° | 0.025 – 0.05 |
| Italy [33] | 38 – 90 | 0.145 | N38-46° | 0.02 – 0.05 |

\* Per 100,000 population. The prevalence of MS for each region is taken from data provided in [42]. A range is given because, often, a range of estimates is available for a particular region.
† Estimates are presented as proband-wise concordance rates [26]. Sometimes concordance was reported as a pair-wise rate and, in these cases, the estimates have been converted into proband-wise rates assuming random sampling of twin-pairs [26]. Nevertheless, in at least some reports [e.g., 32], this assumption is almost certainly violated.
†† For the purposes of determining the probability of “genetic-susceptibility” $\{P(G)\}$ in each region, we have taken: $P(MS) \approx 2 * (prevalence) - see Text$ – and we have adjusted the $MZ$ -twin concordance rates using the Canadian data for differences between fraternal-twin and sibling concordance (*see Text*) as:
Finally, the range of values for $P(G)$ is taken both from the range of the prevalence data and also from the range provided by *Proposition #1* (*see Text*).

Notably, we arrived at this estimate for {*P*(*MS* | *IG*_*MS*_)} by adjusting the observed value of {*P*(*MS* | *MZ*_*MS*_)} downward to account for the presumed impact of the shared *IU* and early post-natal environments of *MZ*-Twins (*see #2 above*). To do this, we estimated this impact from the increased recurrence risks in *DZ*-twins compared to that in non-twin siblings (*see Supplemental Material*). Although, the Canadian data suggests a larger discrepancy between {*P*(*MS*│*DZ*_*MS*_) and *P*(*MS*│*S*_*MS*_)} compared to other studies [27–35], it is still possible that our adjustment is too small. Even so, there is a limit to how large any adjustment can be. Thus, from *Table 2*, it must be the case that:

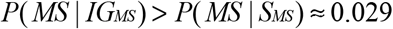

Otherwise, there would be no increased risk of MS in persons who both have 100% of their genes in common and don’t share their *IU* and early post-natal environments compared to persons who both have only 50% of their genes in common and also don’t share their *IU* and early post-natal environments. Importantly, however, even with such an extreme adjustment:

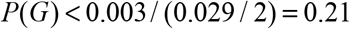

Therefore, even using this extreme estimate, the large majority of the population (>79%) would have no chance of getting MS, regardless of their environmental exposures (*see Proposition #1*).

Nevertheless, these considerations pertain only to the Upper Solution and, as discussed in *#5* (*below*), the observations from Canada regarding recurrence risks for the gender partition in MS make it clear that the set {*X*} is bimodal and, moreover, that it conforms to the Lower Solution. This circumstance will increase the upper limit for genetic susceptibility to MS from the 4.5% estimated here. Nevertheless, even in this circumstance, there are constraints on possible solutions such that:

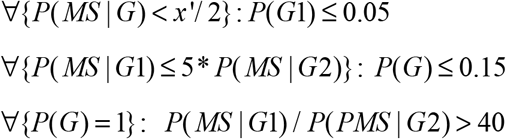

and:

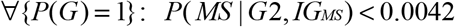

Consequently, although Lower Solutions exist for which, {*P*(*G*) = 1}, none of these simultaneously match the constraints placed the observed the values of (*x*′, *x*_1_′, & *x*_2_′) for the partitions based on gender or *HLA*-status (*see #5 & #6, below*). We conclude, therefore, that the circumstance of {*P*(*G*) = 1} is excluded and that, as a result, developing MS is not a possibility for some portion of the population. Moreover, in earlier iterations of this analysis [3,4,49,50], we defined the subset-(*G*) differently – i.e., as ∀*G*_*i*_ ∈*G* : *P*(*MS* | *G*_*i*_) ≥ *P*(*MS*). It is noteworthy that, in the present analysis, using our new defnitions, our older definition (in the circumstance of the Lower Solution) corresponds to defining only members of the (*G*1)-subset as being “genetically susceptible” to MS (*see Supplemental Material*).

### 5. Genetic Susceptibility in the Gender Partition – *P*(*F│G*) & *P*(*M│G*)

***Conclusions:***

1. The set {*X*} has a bimodal distribution
2. 0.163 ≤ *P*(*MS* | *F*, *G*) ≤ 0.187
3. 0.029 ≤ *P*(*MS* | *M*, *G*) ≤ 0.034
4. 0.23 ≤ *P*(*F* | *G*) ≤ 0.28
5. 4.8 ≤ *P*(*MS* | *F*, *G*) / *P*(*MS* | *M*, *G*) ≤ 6.4
6. 0.041 ≤ *P*(*G*) < 0.047

***Argument:*** For ease of notation, we will use the parameters (*x*, *x*′, *x*_1_, *x*_1_′, *x*_2_, *& x*_2_′) already defined in *Proposition #1* (*above*). The set {*X*} of penetrance values for members of the (*G*)-subset (*Proposition #1*; *Main Text*) is clearly bimodal. Thus, from the data in *Table #2* (*Main Text*):

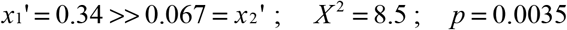

Because we are only considering the possibility of either unimodal or bimodal distributions for the set {*X*}, therefore, the sets (*G*1) and (*G*2), considered separately, must each be unimodal and, thus, meet conditions for the Upper Solution. Using logic identical to that of *Proposition #1* for the (*M / F*) partition, considering susceptible women{(*G*1) = (*F*, *G*)} and susceptible men {(*G*2) = (*M*, *G*)}, using the estimated adjustments for the similar early environment of twins for these two subgroups (*see Supplemental Material*), and using the data provided in *Table 2*, it follows that:

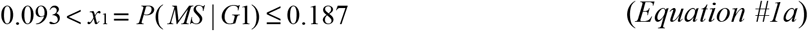

and:

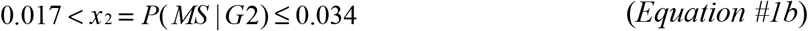

The proportion of MS patients who are women from *Table 2* is 66%. For the WTCCC data this number is 72%. From the study of Orton and colleagues [54] out of Canada, in the most recent epoch, the percentage of MS patients who are women is 76%. Therefore, assuming that: *P*(*F*) = *P*(*M*) = 0.5, and using the smallest (i.e., the most conservative) of these estimated gender imbalances, using the above ranges for men and women, and from the definition of the (*G*)-subset, we can estimate that:

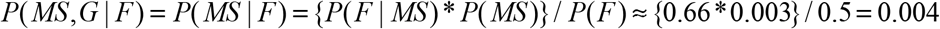

Because:

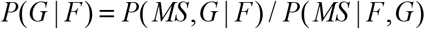

Therefore:

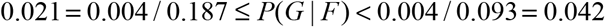

And similarly:

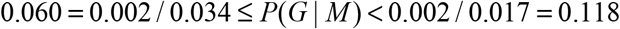

These possible ranges for men and women don’t overlap. Therefore, at a minimum, the excess of men in the (*G*)-subset must be:

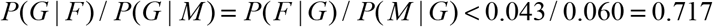

where:

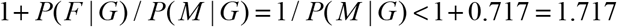

so that:

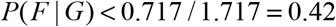

In fact, this gender imbalance is likely even greater than this (*see Supplemental Material*). Thus, there are four serious concerns about undertaking the above calculation. First, in using the above limits for (*x*_1_ and: *x*_2_ – *from Equations #1a & #1b*), we are positing and an extreme and tri-modal distribution for the set {*X*} – i.e., not the unimodal or bimodal distributions under consideration in this manuscript. Thus, this calculation, envisions a distribution, in which half of the women have a penetrance of slightly greater than zero and the other half have a uniform penetrance of (*x*_1_′) – i.e. women have the maximum variance possible – and, in which every man has exactly the same penetrance of (*x*_2_′), which is intermediate between these two extreme penetrance groups of women – i.e. men have a zero variance.

Second, such an extreme distribution seems very unlikely, especially in the circumstance where partitioning the (*G*)-subset by a different MS-associated characteristic – i.e., *HLA-*status (*see #6, below*) – doesn’t even give a hint of the bimodal nature of {*X*}.

Third, it is not possible that the variance of penetrance values for the (*F*, *G*) subset to be at its maximum value. Thus, because (*x*_1_′ > *x*′), then the maximum variance for the (*F*, *G*) -subset – (*x*_1_′/ 2)^2^ – exceeds the maximum total variance possible for the entire (*G*)-subset – (*x*′/ 2)^2^. Consequently, the lower limit for the value of (*x*_1_) in *Equation #1a* – i.e., at its maximum possible variance – must be too low.

And fourth, some of the maximum possible variance in the {*X*} set must be accounted for just by the separation of (*x*_1_) from (*x*_2_) – *see Supplemental Material*. Thus, following the standard development of variance relationships [55], and taking each of these factors into account, leads to the conclusion that:

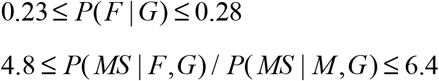

and that:

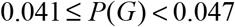

The estimate derived from *Table 2* for the quantity {*P*(*MS* | *M*, *IG*_*MS*_)}, because it is based on only two observations, seems likely to be the least reliable of any in the *Table.* However, even if this estimated penetrance were doubled, there would still be an excess of men in the (*G*)-subset such that:

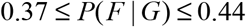

and also:

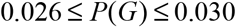

Consequently, not only is genetic susceptibility extremely rare in the population, but also men are more likely than women to be genetically susceptible to MS. At first pass, it might seem biologically improbable that men would be two to four times more likely than women to be in the genetically susceptible subset-(*G*). Thus, if membership in the (*G*)-subset is envisioned as being due to an individual possessing a sufficient combination of some number of loci in a “susceptible state” [56], it is unclear how men could be more likely than women (or *vice versa*) to possess certain combinations and not others. This seems especially unlikely for circumstances, when one association study, specifically focused on the *X*- chromosome, failed to identify any susceptibility loci on this chromosome [7], when another large GWAS found that all but one of the ~200 MS-associated loci were located on autosomal chromosomes [14], and when no major gender interaction term has been reported in the literature. Indeed, considering the different “risk” haplotypes in the *HLA* region identified in the WTCCC, men and women seem equally likely to be carriers [57]. Nevertheless, we can designate (*G_ak_*) to represent each of the (*n*) autosomal genotypes (*k* = 1,2,…*n*) in the general population (*Z*) – i.e., omitting any specification of gender. In this circumstance, it is entirely possible that:

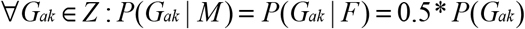

and, yet, for some specific autosomal genotypes to have the characteristic that:

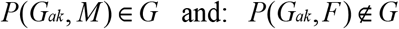

Indeed, such an explanation for the excess in susceptible men would fit well with the observation that the specific genetic combinations, which underlie susceptibility to MS, seem to be unique to each individual (*see #9, below; see also Supplemental Material*). In addition, such a circumstance might also help to rationalize the finding that men likely have a lower threshold of environmental exposure for developing MS compared to women (see *#7, below*).

### 6. Genetic Susceptibility in the HLA Partition – *P*(*G*│*H*+) & *P*(*G*│*H*−)

***Conclusions:***

*P*(*G | H*+) ≈ 3.35* *P*(*G | H*−)
*P*(*G | H*+) ≤ *P*(*G*) / *P*(*H*+) < 0.20

***Argument:*** We will designate individuals who possess 1 or 2 copies of the Class II *HLA-DRB1*15:01~HLA*-*DQB1*06:02~a1* haplotype – i.e. the (*H+*) haplotype – as being members of the (*H+*)-subset and those who possess 0 copies of this haplotype as being members of the (*H−*)-subset. Some epidemiological studies only report *HLA-DRB1*15* or *HLA~DRB1*15:01* carrier status. Nevertheless, because, in the WTCCC, 93.4% of *HLA*-*DRB1*15*-alleles are actually the *HLA-DRB1*15:01* allele, and because 99% of *HLA*-*DRB1*15:01* carriers also carry the full haplotype [58], each of these designations will be used interchangeably as (*H+*).

It is clear that (*H+*) status is considerably enriched in the MS population compared to controls. For example, in WTCCC controls{*P*(*H*+) = 0.23}, whereas in cases{*P*(*H*+ | *MS*) = 0.50}. This enrichment of (*H+*) status in MS could occur in two ways. First, (*H+*) could make membership in the (*G*)-subset more likely than it is for the (*H−*)-subset – i.e., it is due to an impact on the ratio of: *P*(*G* | *H*+) / *P*(*G* | *H*−). Second, members of the (*G*, *H*+) subset may have a greater penetrance for MS than members of the (*G*, *H*−) subset – i.e., it is due to an impact on the ratio of: *P*(*MS* | *G*, *H*+) / *P*(*MS* | *G*, *H*−). The available epidemiological data (*see Supplemental Material*) suggests that the majority of this enrichment is the result of the 1^st^ of these two possible mechanisms and that:

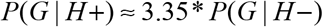

In addition, the observation that only 4.7% (or less) of the population is genetically susceptible, together with the WTCCC observation that: *P*(*H*+) = 0.23, indicates that fewer than 20% (4.7/23) of (*H+*)-carriers are even genetically susceptible to MS. Indeed, taken together, the fact that only half of MS-patients are in the (*H+*) subset, and the fact that this estimate for genetic susceptibility represents an upper bound, indicates that the actual percentage of (*H+*) carriers who are genetically susceptible must be far less than this 20% figure. Nevertheless, essentially all of the conserved extended haplotypes (*CEHs*) that carry (*H+*) – even those unique in the WTCCC dataset – are associated with MS [58]. Therefore, it is likely that the (*H+*)-haplotype, itself, contributes to genetic susceptibility. Despite this contribution, however, the majority of (*H+*)-subset members have no chance whatsoever of developing MS. Therefore, at least with respect to the (*H+*) haplotype, genetic susceptibility to MS must result from the combined effect of (*H+*) together with the effects of certain other (as yet, unidentified) genetic factors (*see Supplemental Material*). By itself, (*H+*) membership poses no MS-risk.

### 7. Environmental Factors in MS

***Conclusions:***

1. Environmental factors are critical to MS pathogenesis
2. Susceptible women are more responsive than men to changes in the environmental conditions related to MS pathogenesis.
3. Compared to women, men have a lower threshold of environmental exposure at which they can develop MS
4. Currently, the environmental factors involved in MS pathogenesis are population-wide exposures.
5. Stochastic factors play an important role in MS pathogenesis

***Argument:*** We define (*E*_*T*_) to be the prevailing environmental conditions (whatever these are) experienced by the population during some time-period (*T*). As noted in the Methods, we define (*E*_*i*_) to be the specific environmental exposure, which is sufficient for MS to develop in the *i*^*th*^ susceptible individual (however many events are involved, whenever these events need to act, and whatever these events might be) – i.e., both the events (*E*_*i*_ and *G*_*i*_) need to occur jointly in order for MS to develop in the (*i*^*th*^) individual. Because genetic susceptibility is independent of the environmental conditions, the probability of a sufficient environmental exposure {*P*(*E*)} in the (*G*)-subset at time-period (*T*) can be expressed as:

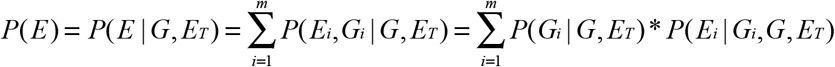

where:

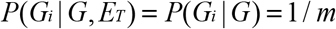

When {*P*(*E*) = 0}, it is not possible for any susceptible person to experience an environment sufficient to cause MS. By contrast, when {*P*(*E*) = 1}, every susceptible person experiences an environment sufficient to cause MS. If some susceptible individuals can develop MS, regardless of their environmental experiences, then: 0 < *P*(*E*) ≤ 1. Importantly, those circumstances, in which {*P*(*E*) = 0}, only imply that, whatever environmental exposures take place (i.e., *E*_*T*_), these are insufficient to cause MS in <u>anyone</u>.

Notably, also, this expression for *P*(*E*) explicitly incorporates the possibility that each genotype in (*G*) may require a unique set of environmental events in order for MS to develop in that individual. Nevertheless, despite this possibility, the existing epidemiological data suggests that many (or most) MS patients are responding to similar environmental events and, thus, any large variability in this regard is probably not a major factor in MS pathogenesis.

For example, despite the fact that every MS patient (except *MZ-*twins) has a unique combination of “states” at the (*>200*) susceptibility loci (*see Supplemental Material*), the data from Canada indicates that the change in general environmental conditions (whatever these are), which have taken place between the time periods of (1941-1945) and (1976-1980), have produced, at a minimum, a 32% increase in the prevalence of MS (*see Supplemental Material*). Moreover, because this increase has occurred world-wide and predominantly in women [3,4,49,50,54], the (*F:M*) sex ratio for MS in Canada has increased during every 5-year increment except one between these two time-periods [54]. Over the entire interval, the ratio has increased from 2.2 in (1941-1945) to 3.2 in (1976-1980). These changes are far too rapid to be genetically based.

It is conceivable that this observed sex-ratio change might be artifactual. For example, if women were more likely than men to have minimally symptomatic MS, then, with such patients now being diagnosed by our improved imaging and laboratory methods, women might represent a disproportionate number of these newly diagnosed cases. Alternatively, in earlier eras, vague symptoms of MS in women may have been written off as “non-organic” more often than they were in men. Nevertheless, four lines of evidence argue strongly against this change being an artifact. First, this increase in the sex ratio began before, and continued up to, the advent of modern imaging and laboratory methods [54]. Second, among asymptomatic individuals, incidentally found to have MS by MRI, the (*F:M*) ratio is approximately the same as current estimates for symptomatic MS and 80% of the those with spinal cord lesions are women – i.e., those lesions having, by far, the greatest odds of any for progression to “clinical” MS [59]. Third, if (as seems likely), women have a higher threshold for developing MS than men, this would require the difference in exposure between the genders to be one of degree not one of kind (see *below; see also Supplemental Material*). Finally, and most persuasively, the greater penetrance of MS in women is confirmed independently by the *MZ*-twin data (*see #5 above*). Consequently, the increase observed in the (*F:M*) sex ratio of Canada [54] must have an environmental basis.

In addition, a prior Epstein Barr viral (*EBV*) infection seems to be a prerequisite for most (or all) genotypes in (*G*) to develop MS [3,4,49,50,60–62]. Indeed, if (as suggested by these studies) a prior *EBV* infection occurs in 100% of MS cases, this would indicate that *EBV* exposure is part of the causal pathway leading to MS and that, at least, this environmental exposure is required for disease pathogenesis [49]. In addition, the likelihood that members of the (*G*)-subset will develop MS seems to be influenced greatly by vitamin D deficiency, latitude, migration, and the *IU* environment [3,4,49,50,60–62]. Each of these additional observations also indicates that similar environmental changes can affect a large proportion of genetically susceptible individuals in a similar manner (i.e., contribute to MS pathogenesis).

Using the standard methods of survival analysis [63], we can define the cumulative survival {*S(u)*} and failure {*F(u)*} functions as well as the hazard-rate functions {*h(u)*} and {*g(u)*} for developing MS at different environmental exposures in “susceptible” men and women (respectively). These hazard-rate functions are assumed (initially) to be proportional. The implications of non-proportionality are considered in the *Supplemental Material* and in the legend of *Fig 3*. However, assuming proportionality, then:

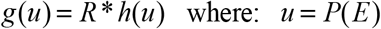

**Figure 3.**
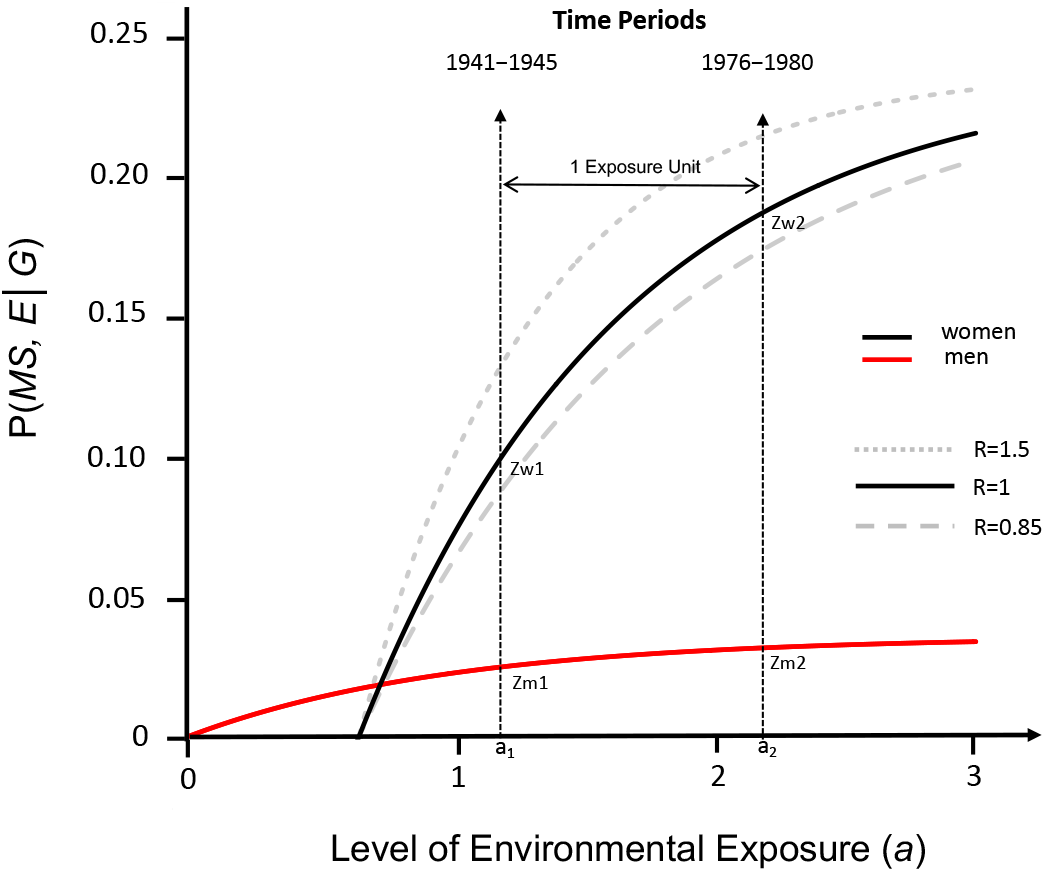
Response curves for the likelihood of developing MS in “genetically susceptible” men and women with an increasing probability of a sufficient environmental exposure{*P*(*E*)}, assuming proportional hazards (*see Text*). Response curves are derived from the change in the (*F:M*) sex-ratio over time in Canada [58] and using the estimates for *P*(*MS*) and *P*(*G*) derived in the *Text*. The probability of getting MS in a “genetically susceptible” individual – i.e., *P*(*MS*, *E* | *G*) – is shown on the y-axis. The exposure level {*P*(*E*)} for the population is shown on the *x-axis* using transformed “exposure units” (*a*) – *see #7, in Text*. Labels for points *Zw* = *P*(*MS*, *E* | *G*, *F*) and *Zm* = *P*(*MS*, *E* | *G*, *M*) are provided at time-points (*1*) and (*2*). One “exposure unit” is defined arbitrarily as: (*a*_2_ − *a*_1_) for men and (*a*_2_ *^app^* − *a*_1_*^app^*) for women (*see Text*). Solid-line plots have been constructed using the known values of (*Zw*_2_) and (*Zm*_2_), together with the conditions: {*C* = 0.6}, {*R* = 1}, {*P*(*G*) = 0.045} and {*P*(*F* | *G*) = 0.28}, the latter being at the top of its estimated range (*see #8 above*). The limiting values for (*Zm*) and (*Zw*) are: (*c* = 0.037) and (*d* = 0.239). Response curves for women under conditions (*R* = 0.85) and: (*R* = 1.5) are also depicted and are shown in grey lines (dashed and dotted, respectively). Changes to the value of (*C*) will slightly alter the units of the y-axis. As seen in the *Figure*, men have a lower threshold for developing MS compared to women (*see Text*), and changes to the value of (*R*) alter how quickly the curves reach their plateau (limit). Increasing the estimate of{*P*(*F* | *G*)} makes the two curves slightly closer to each other. If the hazard is not proportional, for women, each of the points (*Zw*_1_, *Zw*_2_, *Zm*_1_, and *Zm*_2_) would be the same as depicted for (*R* = 1), although the scale of the *x-axis* for the two exponential curves would be transformed non-linearly and, thus, the response-curve in men could not be compared directly to the curve in women. Moreover, the *x-intercept* for the curve in women would be at (*a*^*app*^ = 0). Nevertheless, the limiting values (***c***) and (***d***) would be unchanged and, under any circumstances, women (relative to men) would still be seen to exhibit a greater responsiveness to those changes in environmental exposure, which have taken place between the two time-periods.

For men, we can transform exposure from (*u*) units into (*a*) units, first by defining {*H(u)*}to be the definite integral of the hazard-function {*h(u)*} from a (*u*) level of exposure to a (*0*) level of exposure and, second, by defining the (*a*) units to be:

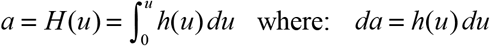

Because these (*a*) units are arbitrary, we can assign “1 unit” of environmental exposure in men to be the difference in exposure level between any two time points (e.g., *a_1_* and *a_2_*) such that:

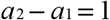

For women, we can similarly transform exposure into a different scale of so-called “apparent” exposure units *(a^app^*) such that:

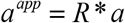

and where we now define “1 unit” of environmental exposure (on this scale) as:

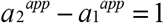

The choice of which gender (*men* or *women*) to assign to which scale is completely arbitrary.

A standard derivation from survival analysis methods [63], demonstrates that the survival curves are exponential with respect to their hazard functions.

Thus, for men:

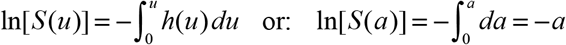

and, for women:

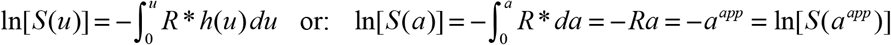

So that, for men:

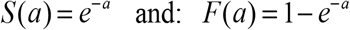

and, for women:

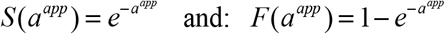

In considering the probability of failure (i.e., of developing MS), we will use subscripts (*1*) and (*2*) to denote the failure probabilities and the values of other parameters at the 1^st^ and 2^nd^ time-periods respectively. Importantly, unlike true survival (where everyone fails given a sufficient amount of time), the probability of developing MS may not become 100% as the probability of a sufficient environmental exposure increases to {*P*(*E*) = 1}. Moreover, the limiting value for the cumulative probability of developing MS in men (***c***) need not be the same as that in women (***d***). However, because the new definition of the subset-(*G*) differs from earlier iterations of our analysis [3,4,49,50], the environmental exposure at which the development of MS becomes possible (i.e., the threshold) must occur at {*P*(*E*) = 0} for, at least, one of the two gender subgroups – provided that this exposure level is possible (*see above*).

From these definitions, the cumulative failure probability for susceptible women (*Zw*) and men (*Zm*) at the 1^st^ time period is:

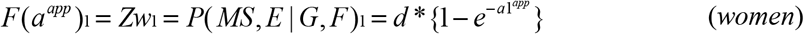

and:

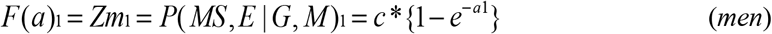

By our definitions of “1 exposure unit”, these equations, at the 2^nd^ time point, become:

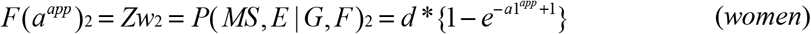

and:

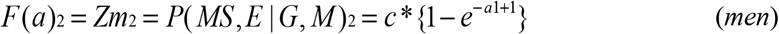

Because the observations at time-periods (*1*) and (*2*) represent two points on the exponential response curves for both women and men, and because any two points on an exponential curve defines the curve (both uniquely and completely), we can use the observations regarding the (*F:M*) sex-ratio change over time in Canada [54], to construct these two response curves.

From the definition of {*P*(*E*)} and using: *P*(*G*, *F*) ≈ 0.28 * *P*(*G*) = 0.013 – i.e., at the top of its estimated range (*see #5 above; see also Supplemental Material*) – and: *P*(*G*) ≈ 0.045 – i.e., in the middle of its estimated range (*see #5 above*) – we can estimate the values of *(Zw_2_*) and *(Zm_2_*) as:

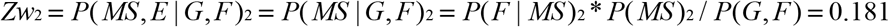

and:

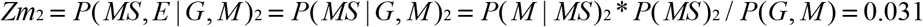

Moreover, as demonstrated in the *Supplemental Material*, assuming that MS prevalence has either remained stable or increased over the time-interval, we can define the term (*C*) such that:

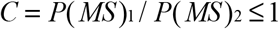

and, thereby, re-express *(Zw_1_*) and *(Zm_1_*) in terms of *(Zw_2_*) and *(Zm_2_*).

Thus:

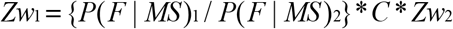

and:

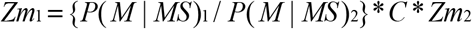

where:

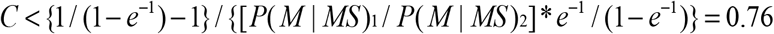

And, thus:

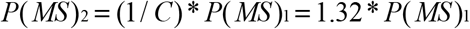

Consequently, based on the population data from Canada, the prevalence of MS must have increased by more than 32% between these two time periods.

Finally (*see Supplemental Material*), we can estimate the value of both (***c***) and (***d***) as:

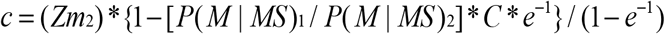

and:

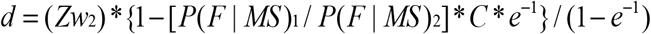

Thus, using the observed change in the (*F:M*) sex-ratio over time in Canada, together with our estimates for *P*(*G*) and *P*(*F│G*), we have all the data needed to construct the complete response curves for the cumulative probability of developing MS in genetically susceptible women and men (*Fig. 3*). What these curves make clear is that both *P*(*E*) and *P*(*MS*) are changing over time, which indicates that specific environmental conditions, in addition to specific susceptible genetic combinations, are necessary for MS to develop. Thus, MS develops when the right genetic constitution is exposed to the right environmental conditions (i.e., it is fundamentally due to a gene-environment interaction).

Because, as noted above, the scales for the response curves in women and men are assumed to be proportional, the plot for *women* on the (*a*) scale will be stretched or compressed (along the *x-axis*) – depending upon the value of (*R*) – compared to the plot for *women* on the (*a*^*app*^) scale (*Fig. 3*). The threshold for men (*λm*) is at the intersection-point {(*a*, *Zm*) = (*λm*,0)} and the threshold for women (*λw*) is at the intersection-point {(*a*, *Zw*) = (*λw*,0)}. By the definitions of (*E*) and (*a*), one of these two thresholds must occur at {(*a*, *Z*) = (0,0)}. However, these thresholds need not be the same, so we define the threshold difference between women and men as: (*λ* = *λw*− *λm*). Thus, if women have a higher threshold than men: (*λ* > 0). On the (*a*^*app*^) scale, the *x-intercept* always occurs at {*a*^*app*^ = *λw* = 0}, and, thus, (*λw*) is independent of *R*.

Three final points are also worth making. First, because (*λw*) is independent of *R*, we can use the condition of (*R* = 1) to evaluate (*λw*). In this circumstance, these exponential equations (*see Supplemental Material*) can be re-arranged to yield:

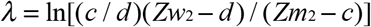

Consequently, basic epidemiologic data can be used to determine the difference in threshold (*λ*) that exists between women and men. This analysis leads to the conclusion that:

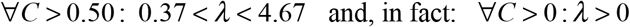

Therefore, if the hazards are proportional, men have a lower threshold for developing MS compared to women (*Fig.3; Supplemental Material*). A lower threshold in men is also suggested by a report from Europe and the United States [64], which found that, prior 1922, men accounted for 58% of the MS cases (*Table 4*). By our definition of *P*(*E*) these thresholds indicate the exposure, at which MS becomes possible. If women required a fundamentally different kind of exposure than men, it would be very hard to rationalize a difference in threshold because, in such a circumstance, in some environments, women would be more likely and, in other environments, less likely than men to receive the correct exposure. Rather, a difference in threshold implies that men and women are responding to similar events but that men require a less extreme degree of exposure in order to develop MS. For example, perhaps, men become susceptible with a lesser degree of vitamin D deficiency or with EBV infection occurring over a broader age-range compared to women.

**Table 4.** Sex Distribution of Multiple Sclerosis cases reported prior to 1922*

|  | **Europe**<br>**(Historical)** | **United States**<br>**(Historical)** | **United States**<br>**(Wechsler Series)** | **Total** |
| --- | --- | --- | --- | --- |
| **Men** | 658 (58%) | 99 (60%) | 117 (59%) | 874 (58%) |
| **Women** | 484 (42%) | 67 (40%) | 80 (41%) | 631 (42%) |
| **Total** | 1,142 (100%) | 166 (100%) | 197 (100%) | 1,505 (100%) |

\* Data from: Wechsler IS [64]. Historical cases were reported in the medical literature, for the most part, prior to 1903, whereas Wechsler's series was drawn from the Mount Sinai Hospital, the Montefiore Hospital, and the Vanderbilt Clinic (1912-1921).

Second, we note that: {*P*(*MS* | *G*, *E*, *M*) = *c*}, so that (*Zm*_2_) can be re-expressed as:

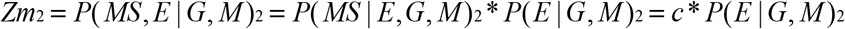

This equation can be rearranged to yield:

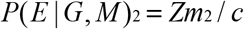

From above:

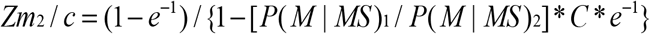

Assuming that there has been less than a two-fold increase in MS prevalence in Canada over the 35-year interval (i.e., *C* > 0.5) then:

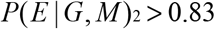

And, similarly, for women:

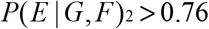

These results strongly suggest that the relevant environmental exposures (especially when these are multiple) are currently occurring at population-wide levels. For example, if three, equally likely and independent, environmental events (*EE_1_, EE_2_,* and *EE_3_*) – possibly sequential [49,50] – were necessary to produce MS in a susceptible individual, then:

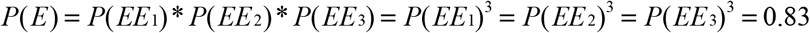

or:

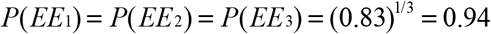

so that, under the stated circumstances, more than 94% of the population would experience each environmental event. Such a conclusion is fully consistent with the same conclusion reached from studies in adopted individuals, in siblings and half-siblings raised together or apart, in conjugal couples, and in brothers and sisters of different birth order, which have generally indicated that MS-risk is currently unaffected by the micro-environment of families [65–71].

And third, it is clear that both of these response curves plateau well below 100% failure (*Fig 3*). Therefore, there must be stochastic processes that partially determine whether a susceptible individual with a sufficient environmental exposure will actually develop disease (*see #9, below*).

### 8. The Future Fate of *P*(*MS│IG_MS_*)

***Conclusions:***

1. 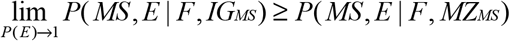
2. 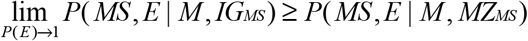

***Argument:*** As a sufficient environmental exposure {*P*(*E*)} becomes more likely, the quantity {*P*(*MS│IG_MS_*)} will, of necessity, change. Earlier, we described this term as having removed the impact of the shared *IU* and early post-natal environments of *MZ*-twins. This description, however, is not quite accurate. For example, we can break down a “sufficient” environmental exposure (*see Supplemental Material*) into those factors that are shared exclusively by *MZ*-twins (*E*_1_), those factors that are shared by the population generally (*E*_2_), and those factors that shared exclusively within the family microenvironment (*E*_3_). As noted above, however, the family microenvironment seems not to have any impact on the likelihood of MS [65–71]. In this circumstance, because only factors (*E*_1_ and *E*_2_) are necessary for a sufficient exposure, then:

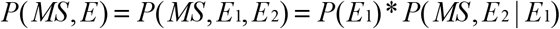

and:

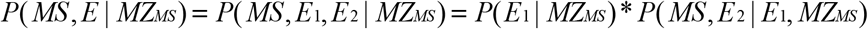

If an individual’s identical twin is known to have MS, it is likely that this individual, also, has experienced a “sufficient” *IU* and early post-natal environment.

Conceived of in this way, the term {*P*(*MS│IG_MS_*)} can be rewritten as:

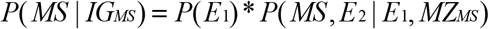

and the adjusted penetrance {*P*(*MS│IG_MS_*)} hasn’t really “removed” the impact of these early environmental similarities. Rather, {*P*(*E*_1_ | *MZ*_*MS*_)} has simply been reset to its population level {*P*(*E*_1_)}. Because *MZ-*twins share both identical genotypes and the same *IU* and early post-natal environment, we expect that: *P*(*E*_1_ | *MZ*_*MS*_) = 1. Consequently, as {*P*(*E*_1_)} increases in the population to:

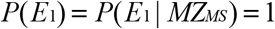

the term {*P*(*MS│IG_MS_*)} will approach, and ultimately reach:

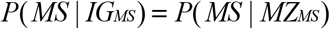

In this case, therefore, the limiting value for {*P*(*MS*, *E* | *G*)} in men (***c***) and women (***d***) – *see #7 above* – must conform to the constraints of:

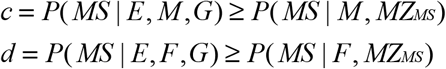

The reason for the inequality is that, in those circumstance where:

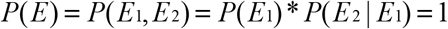

it must be that both: *P*(*E*_1_) = 1 and: *P*(*E*_2_ | *E*_1_) = 1. Naturally, the fact that {*P*(*E*_1_)} has increased to unity does not guarantee that {*P*(*E*_2_ | *E*_1_)} has done the same, so that the limiting value for {*P*(*MS*, *E* | *G*) = *P*(*MS* | *G*)} may be greater than {*P*(*MS* | *G*, *MZ*_*MS*_) = *P*(*MS* | *MZ*_*MS*_)}.

Nevertheless, if it is currently true (*see #7 above*), that:

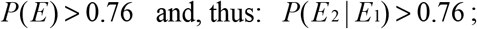

then it must also true that:

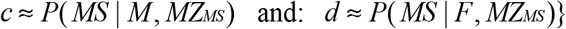

Regardless, however, the depicted curves (*Fig. 3*) must be inaccurate because, in the Figure:

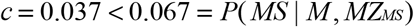

and:

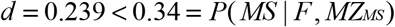

Clearly there are several variables that can be adjusted {*C*, *R*, *P*(*G*), *P*(*F* | *G*), and *P*(*MS*)} to match the values for both (***c***) and (***d***) with these observed *MZ*-twin concordance rates. Therefore, it is possible to consider different combinations of these variables and determine those combinations that match these constraints. For example, using variables ranges of: (0.5 ≤ *C* ≤ 0.75), (0.25 ≤ *R* ≤ 4.5), (0.001 ≤ *P*(*G*) < 0.10), (0.23 ≤ *P*(*F* | *G*) ≤ 0.70), and (0.0025 ≤ *P*(*MS*) ≤ 0.006), and further requiring that the estimates for (***c***) and (***d***) to be within (± 15%) of the observed values for the proband-wise *MZ*-twin concordance rates (*Table 2*), there are numerous combinations, which match these constraints. The solution space covered by these combinations includes the full range of possibilities for the parameters of (*C*) and (*R*). By contrast, the ranges for both *P*(*F|G*) and *P*(*G*) are restricted: {0.35 ≤ *P*(*F* | *G*) ≤ 0.5} and {0.02 ≤ *P*(*G*) ≤ 0.05}. This restricted range for *P*(*G*) fits, generally, within the framework developed previously (*see #5 above*). The range for *P*(*F|G*), however, is outside of the range developed previously (*see #5 above*) although, as discussed in the *Supplemental Material*, this could relate to an underestimate for the parameter {*P*(*MS* | *M*, *IG*_*MS*_)} from *Table 2*.

Also, 94% of these potential solutions require the condition that {*P*(*MS*) > 0.003}. This latter circumstance might reflect an under-ascertainment of cases when estimating the disease prevalence for the general population (*Z*). Indeed, several autopsy studies have indicated that the prevalence of undiagnosed (pathological) MS is ~0.1% [45–48]. Thus, with minimally symptomatic (or asymptomatic) MS occurring in as many 0.1% of the population, this could potentially increase the estimated *P*(*MS*) by as much as 50–100%. Although, such diagnostic errors are probably less common in the modern era, many minimally symptomatic (or asymptomatic) patients are still being undiagnosed during life [59]. Moreover, any such under-ascertainment is likely to be less for *MZ*-twins, *DZ*-twins, and siblings. For example, an initially unaffected twin or non-twin sibling of a patient with MS will, almost certainly, be more carefully monitored for possible MS symptoms (i.e., for minimally symptomatic presentations) than will an individual in the general population. In such a circumstance, these diagnostic failures will be fewer in the (*MZ*_*MS*_), (*DZ*_*MS*_), and (*S*_*MS*_) populations than in the general population and the *MZ*-twin concordance rates will, thus, provide a more accurate reflection of the maximum likelihood of getting MS {i.e., *P*(*MS* | *G*, *E*_1_, *E*_2_)} than will those estimates of *P*(*MS*) derived from the *MS*-prevalence in the general population. Such a circumstance might help to rationalize this apparent discrepancy.

### 9. Missing Heritability?

***Conclusions:***

1. Both “genetic” and “environmental” factors are necessary for MS expression; Neither alone are sufficient.
2. A large portion of the “causal pathway” to MS is stochastic
3. There is no need to invoke any “missing heritability” in MS

***Argument:*** Only a small proportion of the population seems to be genetically susceptible to developing MS, which implies that this is a “genetic” disorder. In addition, a suitable environmental exposure, like a suitable genetic constitution, is also a necessary part of MS pathogenesis. Despite this, however, the combination of a susceptible genotype together with a sufficient environmental exposure, does not invariably lead to the disease of MS and, in fact, the response curves in both men and women plateau well below 100% (*Fig 3*), even when everyone receives an environmental exposure suitable for their particular genotype – i.e., when {*P*(*E*) = 1}. This variance in the likelihood of getting MS for certain susceptible genotypes cannot be attributed to unidentified environmental conditions because the definition of the term {*P*(*E*)} – *see #7 above* – explicitly includes all such factors, both if they are known (or suspected) and also if they are completely unknown. Therefore, a large portion of the overall variance in MS disease-expression must be due to stochastic processes.

In this context, dividing the total variance in disease expression into genetic and environmental components, at least for MS, mischaracterizes the situation. This has important implications regarding current estimates for the “missing hereditability” in MS [71–73]. First, as noted above, a large portion of the variability in MS expression must be due to stochastic processes that are neither environmental nor genetic. And second, specific gene-gene combinations (likely unique to individuals or very small groups of individuals) must underlie genetic susceptibility to MS (*see #6 above; see also Supplemental Material*). Thus, with over 200 MS-associated loci [14], each (potentially) having more than one “susceptible state” (e.g., the *MHC*), the number of possible combinations of states at these loci is so huge that, almost certainly, everyone (except *MZ*-twins) possesses a unique combination of these “susceptible states” (*see Supplemental Material*). Indeed, considering (*H+*)-status together with only the first 102 of these MS-associated SNPs [13], everyone (including both cases and controls) in the WTCCC population does, in fact, possess a unique combination (*Supplemental Material*). Consequently, if only a few such combinations are members of the (*G*)-subset, even among those combinations that are quite similar to each other (*see #6 above; see also Supplemental Material*), then there are more than enough genetic associations already identified to account fully for (*G*)-subset membership. Naturally, many more loci may yet be identified, although positing their existence is unnecessary.

Alternatively, if “missing heritability” is only meant to imply that our genetic model cannot predict accurately the occurrence of MS, then, indeed, almost all of the heritability of MS remains unexplained. Thus, the environmental factors, the actual (as opposed to associated) genetic factors involved in causing disease, the necessary gene-gene combinations, the various gene-environment interactions, and the stochastic factors – all of which contribute importantly to whether MS can, or will, develop in a specific individual – are poorly understood, thereby making any accurate prediction of MS occurrence impossible at present.

## Discussion

The present analysis provides considerable insight to the nature and basis of susceptibility to MS and to the role of genetic determinants in polygenic diseases. Firstly, we establish that, fundamentally, MS is a genetic disorder. Only a tiny fraction of the population (less than 4.7%) is susceptible. Thus, more than 95.3% of the population has no chance of developing MS, regardless of the environmental conditions that these individuals experience. Thus, the correct genetic make-up is essential for disease pathogenesis. The basis of this genetic susceptibility, however, is complex. Single genes or single haplotypes do not contribute much. For example, in MS, the Class II *HLA-DRB1*15:01~HLA-DQB1*06:02~a1* (*H+*)-haplotype is the genetic trait with the largest (by far) disease-association of any in the genome (for the WTCCC: *OR=*3.28; *p*≪10^−300^). Nevertheless, despite this strong association more (and, likely, far more) than 80% of individuals who carry this haplotype have no MS-risk whatsoever. In this circumstance, it must be that genetic susceptibility depends upon the possession of this haplotype in combination with other genetic traits. Notably, this haplotype is only a part of much larger *CEHs*, which span the entire *MHC* region [23,24]. Even considering the large number and variety of these highly selected *CEHs*, however, genetic susceptibility cannot be explained on the basis of the state of the *MHC*. Despite a significant variability in the observed disease-association among the different (*H+*)-carrying *CEHs*, every such *CEH* (regardless of its rarity) is strongly MS-associated [23,24].

Moreover, it seems clear that, although certain genetic combinations increase the likelihood of (*G*)-subset membership, the actual combinations that do this are quite heterogeneous, and only a small proportion of genetically susceptible individuals (who actually develop MS) share even the same 4-locus genetic combination (*Supplemental Material*). These observations also suggest that susceptibility to MS, although genetically based, is idiosyncratic.

Despite the conclusion that MS is genetic, however, MS is equally an environmental disease. Specific environmental exposures are also necessary for disease-patho*g*enesis. Indeed, the fact that there has been a marked recent increase in both MS-prevalence and the (*F:M*) sex-ratio, indicates that a sufficient environmental exposure is required for MS to develop (*Fig. 3*). If you are not exposed to a sufficient environment, you cannot get MS, regardless of your genetic make-up. However, neither environment nor genetics alone is sufficient. Rather, MS is due to an interaction between the two.

Several environmental events, probably sequential, seem necessary for MS to develop in a genetically susceptible individual [3,4,49,50,60–62]. The first environmental event, as discussed previously [49], is one that occurs during *IU* or early post-natal period. Support for such a factor comes from the discrepancy in recurrence-rates between twin and non-twin siblings, from the fact that concordant half-twins are twice as likely to share the mother than the father, and from the periodic, circa-annum, effect that month-of-birth has on the subsequent likelihood of developing MS [49]. In the northern hemisphere, this periodicity to MS-susceptibility peaks just before the summer months and dips to its nadir just before winter and this pattern is inverted southern hemisphere [49]. Such a pattern of periodicity implicates an environmental factor, occurring near birth, that is coupled to the solar cycle [49].

A second environmental event is implied by the published migration data [49]. Thus, when an individual relocates (prior to ~15 years of age) from an area of high-prevalence to an area of low-prevalence (or *vice versa*), their MS risk is similar to that of the area <u>to which</u> they moved. By contrast, when they make the same relocation after this time, their MS risk seems to remain that of the area <u>from which</u> they moved. These observations implicate an environmental event, involved in MS-pathogenesis, which occurs at or around puberty [49]. And third, the clinical onset of MS generally occurs long after the first and second events have already taken place (*Fig. 1*), suggesting that one or more additional environmental events are also required for clinical MS to develop.

Naturally, there is no guarantee that the environmental events, which are sufficient to cause MS in one person, are the same as those that are sufficient in another. Nevertheless, those factors or events, which have been implicated in MS-pathogenesis so far, appear to affect a large proportion of susceptible individuals in a similar manner. Thus, the fact that we even have evidence for the first two factors (*as described above*) suggests this. In addition, a prior Epstein Barr viral (*EBV*) infection has been strongly linked to MS, especially when this infection results in symptomatic mononucleosis. Indeed, such an infection prior to clinical onset occurs in ~100% of MS cases [3,4,49,50,60–62] and, if this is the case, this would indicate that *EBV* exposure is a necessary part of the causal pathway leading to MS [49]. Finally, there is a considerable amount of circumstantial evidence, which suggests a role for vitamin D deficiency in this causal pathway [49].

However, even when the correct genetic background occurs together with an environmental exposure sufficient to cause MS in someone of that background, more than 50% of such individuals will still not develop clinical disease. Some of these individuals, no doubt, will have subclinical disease [45–48,59]. However, although such a circumstance, will increase our estimate of {*P*(*MS*)} by as much as 50-100%, this is still insufficient to get the plateaus of the response curves (*Fig. 3*) to exceed the 50% mark. In men (who have a plateau significantly lower than that of women), this conclusion is even more evident (*Fig. 3*). Consequently, because a sufficient environmental exposure has been defined broadly (to include both factors that are known or suspected as well as factors that are completely unknown), the fact that some individuals with the proper combination of genes and environment still fail to develop disease, indicates that stochastic processes are also involved in disease-pathogenesis.

And finally, it is worth noting that the nature of genetic susceptibility developed in this manuscript is applicable to a wide range of other complex polygenetic disorders such as type-1 diabetes mellitus, celiac disease, and rheumatoid arthritis. Indeed, based solely upon *Proposition #1*, if the proband-wise *MZ*-twin concordance rate, for any disease, greatly exceeds the prevalence of disease in the general population, then only a tiny fraction of the population has any possibility of getting the illness. Moreover, any disease for which the proband-wise *MZ*-twin concordance rate is substantially less than 100% must, in addition to genetic susceptibility, include environmental factors, stochastic factors, or both in the causal pathway leading to the disease.

## Notes

### Competing Interest Statement

The authors have declared no competing interest.

## References

1. Gourraud PA, Harbo HF, Hauser SL, Baranzini SE. (2012) The genetics of multiple sclerosis: an up-to-date review. Immunol Rev 248:87–103.

2. Hofker MH, Fu J, Wijmenga C. (2014) The genome revolution and its role in understanding complex diseases. Biochim Biophys Acta 1842:1889–1895.

3. Goodin DS. The nature of genetic susceptibility to multiple sclerosis: Constraining the Possibilities. BMC Neurology 2016;16:56.

4. Goodin DS. The Genetic and Environmental Bases of Complex Human-Disease: Extending the Utility of Twin-Studies. PLoS One 2012;7(12): e47875.

5. GAMES, the Transatlantic Multiple Sclerosis Genetics Cooperative. (2003) A meta-analysis of whole genome linkage screens in multiple sclerosis. J Neuroimmunol 2003;143:39–46.

6. de Bakker PIW, Yelensky R, Pe’er I, et al. Efficiency and power in genetic association studies. Nat Genet 2005;37:1217–1223.

7. Herrera BM, Cader MZ, Dyment DA, et al. Multiple sclerosis susceptibility and the X chromosome. Mult Scler 2007;13:856–8.

8. The Wellcome Trust Case Control Consortium & The Australo-Anglo-American Spondylitis Consortium. Associations can of 14,500 nonsynonymous SNPs in four diseases identifies autoimmunity variants. Nature Genet 2007;39:1329–1337.

9. Baranzini SE, Wang J, Gibson RA, et al. Genome-wide association analysis of susceptibility and clinical phenotype in multiple sclerosis. Hum Mol Genet. 2009;18:767–778.

10. De Jager PL, Jia X, Wang J, et al. Meta-analysis of genome scans and replication identify CD6, IRF8 and TNFRSF1A as new multiple sclerosis susceptibility loci. Nature Genet 2009;41:776–782.

11. Sanna, S. Pitzalis M, Zoledziewska M, et al. Variants within the immunoregulatory CBLB gene are associated with multiple sclerosis. Nature Genet 2010;42:495–497.

12. The International Multiple Sclerosis Genetics Consortium & the Wellcome Trust Case Control Consortium. Genetic risk and a primary role for cell-mediated immune mechanisms in multiple sclerosis. Nature 2011;476:214–219.

13. International Multiple Sclerosis Genetics Consortium (IMSGC). Analysis of immune-related loci identifies 48 new susceptibility variants for multiple sclerosis Nat Genet 2014;45:1353–60.

14. International Multiple Sclerosis Genetics Consortium. Multiple sclerosis genomic map implicates peripheral immune cells and microglia in susceptibility. Science 2019;65: eaav7188.

15. Dyment DA, Herrera BM, Cader Z, et al. Complex interactions among MHC haplotypes in multiple sclerosis: susceptibility and resistance. Hum Mol Genet 2005;14:2019–2026.

16. Hafler, DA, Compston A, Sawcer S, et al. Risk alleles for multiple sclerosis identified by a genomewide study. N. Engl. J. Med. 2007;357, 851–862.

17. Ramagopalan, SV, Anderson, C, Sadovnick, AD, Ebers, GC. Genomewide study of multiple sclerosis. N. Engl. J. Med. 2007;357, 2199–2200.

18. Link J, Kockum I, Lorentzen AR, et al. Importance of Human Leukocyte Antigen (HLA) Class I and II Alleles on the Risk of Multiple Sclerosis. PLoS One 2012; 7(5):e36779.

19. Patsopoulos NA, Barcellos LF, Hintzen RQ, et al. (2014) Fine-Mapping the Genetic Association of the Major Histocompatibility Complex in Multiple Sclerosis: HLA and Non-HLA Effects. PLoS Genet 9(11):e1003926.

20. Chao MJ, Barnardo MC, Lincoln MR, Ramagopalan SV, et al. HLA class I alleles tag HLA-DRB1*1501 haplotypes for differential risk in multiple sclerosis susceptibility. Proc Natl Acad Sci USA 2008;105:13069–74.

21. Lincoln MR, Ramagopalan SV, Chao MJ, Herrera BM, et al. Epistasis among HLA-DRB1, HLA-DQA1, and HLA-DQB1 loci determines multiple sclerosis susceptibility. Proc Natl Acad Sci USA 2009;106:7542–7.

22. Multiple Sclerosis Genetics Group. Linkage of the MHC to familial multiple sclerosis suggests genetic heterogeneity. Hum Molec Genet 1998;7:1229–1234.

23. Goodin DS, Khankhanian P. Single Nucleotide Polymorphism (SNP)-Strings: An Alternative Method for Assessing Genetic Associations. PLoS One 2014;9(4):e90034.

24. Khankhanian P, Gourraud PA, Lizee A, Goodin DS. Haplotype-based approach to known MS-associated regions increases the amount of explained risk. J Med Genet. 2015;52:587–594.

25. Harrison’s Principles of Internal Medicine, 18th Edition. Longo DL, Kasper, DL, Jameson JL, Fauci AS, Hauser SL, Loscalzo JL (Eds), McGraw Hill Medical, New York, 2012

26. Witte JS, Carlin JB, Hopper JL. Likelihood-Based Approach to Estimating Twin concordance for dichotomous traits. Genetic Epidemiol. 1999;16:290–304.

27. French Research Group on Multiple Sclerosis. Multiple sclerosis in *54* twinships: Concordance rate is independent of zygosity. Ann Neurol 1992;32:724–727.

28. Mumford CJ, Wood NW, Kellar-Wood H, Thorpe JW, Miller DH, Compston DA. The British Isles survey of multiple sclerosis in twins. Neurology. 1994;44:11–5.

29. Willer CJ, Dyment DA, Rusch NJ, Sadovnick AD, Ebers GC, the Canadian Collaborative Study Group. Twin concordance and sibling recurrence rates in multiple sclerosis. Proc Natl Acad Sci U S A. 2003;100:12877–82.

30. Hansen T, Skytthe A, Stenager E, Petersen HC, Brønnum-Hansen H, Kyvik KO. Concordance for multiple sclerosis in Danish twins: an update of a nationwide study. Mult Scler. 2005;11:504–10.

31. Hansen T, Skytthe A, Stenager E, Petersen HC, Kyvik KO, Brønnum-Hansen H. Risk for multiple sclerosis in dizygotic and monozygotic twins. Mult Scler. 2005;11:500–3.

32. Islam T, Gauderman WJ, Cozen W, et al. Differential twin concordance for multiple sclerosis by latitude of birthplace. Ann Neurol 2006; 60: 56–64.

33. Ristori G, Cannoni S, Stazi MA, et al. and the Italian Study Group on Multiple Sclerosis in Twins. Multiple sclerosis in twins from continental Italy and Sardinia: A Nationwide Study Ann Neurol 2006;59:27–34.

34. Kuusisto H, Kaprio J, Kinnunen E, et al. Concordance and heritability of multiple sclerosis in Finland: Study on a nationwide series of twins. Eur J Neurol 2008;15: 1106–1110.

35. O’Gorman C, Lin R, Stankovich J, Broadley SA. Modeling genetic susceptibility to multiple sclerosis with Family Data. Neuroepidemiology 2013;40:1–12.

36. Liguori M, Marrosu MG, Pugliatti M. et al. Age at onset in multiple sclerosis. Neurol Sci 2000;21:S825–S829.

37. Grytten TN, Lie SA, Aarseth JH, Nyland H, Myhr KM (2008) Survival and cause of death in multiple sclerosis: results from a 50-year follow-up in Western Norway. Mult Scler 2008;14: 1191–1198.

38. Ragonese P, Aridon P, Mazzola MA, Callari G, Palmeri B, et al. Multiple sclerosis survival: a population-based study in Sicily. Eur J Neurol 2010;17: 391–397

39. Kingwell E, van der KM, Zhao Y, Shirani A, Zhu F, et al Relative mortality and survival in multiple sclerosis: findings from British Columbia, Canada. J Neurol Neurosurg Psychiatry 2012;83: 61–66.

40. Scalfari A, Knappertz V, Cutter G, Goodin DS, Ashton R, Ebers GC. Mortality in patients with multiple sclerosis Neurology 2013;81:184–192

41. Goodin DS, Corwin M, Kaufman D, et al. Causes of death among commercially insured multiple sclerosis patients in the United States PLoS One 2014;9(8): e105207.

42. Rosati G. The prevalence of multiple sclerosis in the world: an update. Neurol Sci. 2001;22:117–39.

43. Sundström P, Nyström L, Forsgren L. Incidence (1988-97) and prevalence (1997) of multiple sclerosis in Västerbotten County in northern Sweden. J Neurol Neurosurg Psychiatry. 2003;74:29–32.

44. Harding K, Zhu F, Alotaibi MD, Dugan T, Tremlett H, Kingwell E. Causes that contribute to deaths due to multiple sclerosis: analysis of population-based multiple-cause-death data. Presentation 144. ECTRIMS 2018, Berlin.

45. Vost A, Wolochow D, Howell D. Incidence of infarcts of the brain in heart diseases. J Path Bact 1964;88:463–470.

46. Georgi VW. Multiple Sklerose: Pathologisch-Anatomische Befunde multiple Sklerose bei klinisch nicht diagniostizierte Krankbeiten. Schweiz Med Wochenschr 1966;20:605–607.

47. Gilbert J, Sadler M. Unsuspected multiple sclerosis. Arch Neurol 1983;40:533–536.

48. Engell T. A clinical patho-anatomical study of clinically silent multiple sclerosis. Acta Neurol Scand 1989;79:428–430.

49. Goodin DS. The causal cascade to multiple sclerosis: A model for MS pathogenesis. PLoS One 2009;4(2):e4565.

50. Goodin DS. The epidemiology of multiple sclerosis: Insights to a causal cascade. Handb Clin Neurol. 2016;138:173–206.

51. Hankins GVD, Saade GR. Factors influencing twins and zygosity. Paediatr Perinat Epidemiol 2005 19(Suppl 1):8–9.

52. Jacobson HI. The maximum variance of restricted unimodal distributions. Ann Math Stat. 1969;40:1746–52.

53. Freeman JB, Dale R. Assessing bimodality to detect the presence of a dual cognitive process. Behav Res. 2013;45:83–97.

54. Orton SM, Herrera BM, Yee IM, et al., and the Canadian Collaborative Study Group. (2006) Sex ratio of multiple sclerosis in Canada: A longitudinal study. Lancet Neurol. 5:932–6.

55. Freund JE, Walpole RE. Mathematical Statistics. Prentice Hall, Inc., New Jersey, 1980, pp. 452–473.55.

56. Goodin The genetic basis of multiple sclerosis: a model for MS susceptibility. BMC Neurology 2010, 10:101.

57. Goodin DS, Khankhanian P, Gourraud PA, Vince N. Genetic susceptibility to multiple sclerosis: Interactions between conserved extended haplotypes of the MHC and other susceptibility regions. (*submitted*)

58. Goodin DS, Khankhanian P, Gourraud PA, Vince N. (2018) Highly conserved extended haplotypes of the major histocompatibility complex and their relationship to multiple sclerosis susceptibility. PLoS One 13(2):e0190043.

59. Okuda DT, Mowery EM, Cree BAC, Crabtree EC, Goodin DS, Waubant E, Pelletier D. Asymptomatic spinal cord lesions predict disease progression in radiologically isolated syndrome. Neurology 2011;76:686–692.

60. Ascherio, A., Munger, K.L. Environmental risk factors for multiple sclerosis. Part I: the role of infection. Ann. Neurol. 2007;61, 288–299.

61. Ascherio, A., Munger, K.L.. Environmental risk factors for multiple sclerosis. Part II: noninfectious factors. Ann. Neurol. 2007;61, 504–513.

62. Ascherio, A., Munger, K.L., Simon, K.C., 2010. Vitamin D and multiple sclerosis. Lancet Neurol. 2010;9, 599–612.

63. Fisher LD, van Belle G. Biostatistics: A Methodology for the Health Sciences, John Wiley & Sons, New York, 1993, pp. 786–829.

64. Wechsler IS. Statistics of multiple sclerosis: Including a study of the infantile, congenital, familial, and hereditary forms and the mental and psychic symptoms. Arch Neurol Psychiatr 1922;8:59–75.

65. Bager P, Nielsen NM, Bihrmann K, Frisch M, Wohlfart J, et al. (2006) Sibship characteristics and risk of multiple sclerosis: A nationwide cohort study in Denmark. Am J Epidemiol 163:1112–1117.

66. Compston A, Coles A (2002) Multiple sclerosis. Lancet 359:1221–31.

67. Dyment DA, Yee IML, Ebers GC, Sadovnick AD, and the Canadian Collaborative Study Group (2006) Multiple sclerosis in step siblings: Recurrence risk and ascertainment. J Neurol Neurosurg Psychiatry 77:258–259.

68. Ebers GC, Sadovnick AD, Dyment DA, Yee IM, Willer CJ, et al. (2004) Parent-of-origin effect in multiple sclerosis: observations in half-siblings. Lancet 363:1773–1774.

69. Ebers GC, Yee IML, Sadovnick AD, Duquette P, and the Canadian Collaborative Study Group (2000) Conjugal multiple sclerosis: Population based prevalence and recurrence risks in offspring. Ann Neurol 48:927–931.

70. Sadovnick AD, Yee IML, Ebers GC, and the Canadian Collaborative Study Group (2005) Multiple sclerosis and birth order: A longitudinal cohort study. Lancet Neurol 4:611–617.

71. Sadovnick AD, Ebers GC, Dyment DA, Risch NJ, and the Canadian Collaborative Study Group (1996) Evidence for genetic basis of multiple sclerosis. Lancet 347:1728–1730.

71. Zuk O, Hechter E, Sunyaev SR, Lander ES. The mystery of missing heritability: Genetic interactions create phantom heritability. Proc Natl Acad Sci U S A. 2012;109:1193–1198.

72. Lill CM. Recent advances and future challenges in the genetics of multiple sclerosis. Front Neurol 2014;5:00130 eCollection.

73. Bashinskaya VV, Kulakova OG, Boyko AN, Favorov AV, Favorova OO A review of genome-wide association studies for multiple sclerosis: classical and hypothesis-driven approaches. Hum Genet. 2015;134:1143–62.

