## Supplemental Material for "The Nature of Genetic Susceptibility to Multiple Sclerosis"

   *6e. Proportional vs. Non-proportional Hazard-rates for Men and Women....................…… p.16*

7. *Uniqueness of Susceptible Genotypes in the Population…………….………….......................… p.17*

### 1. Penetrance Adjustment for the Shared Early Environment of Twins

***Conclusions:*** 1. $P(MS \mid IG_{MS}) = \{P(MS \mid S_{MS}) / P(MS \mid DZ_{MS})\} * P(MS \mid MZ_{MS})$

***Argument:*** We define a necessary and sufficient environmental exposure, $P(E)$, to be:

$$P(E) = P(E_1, E_2, E_3) = P(E_1, E_2) \quad \text{where:} \quad P(MS, E) = P(MS)$$

Here ($E_1$) represents that part of this exposure shared exclusively by twins, ($E_2$) represents that part shared by the population generally. By contrast, ($E_3$), the potential impact of the microenvironment exclusively shared by families, seems to have little or no impact on the likelihood of MS [65-71]. We also define $\{P(MS \mid IG_{MS})\}$ to represent the *MZ*-twin concordance rate adjusted to exclude the impact of twins sharing their *IU* and early post-natal environments. This relationship can be stated (*see Main Text*) as:

$$P(MS \mid MZ_{MS}) = P(MS, E_1, E_2 \mid MZ_{MS}) = P(E_1 \mid MZ_{MS}) * P(MS, E_2 \mid E_1, MZ_{MS})$$

Where: $P(MS \mid IG_{MS}) = P(E_1) * P(MS, E_2 \mid E_1, MZ_{MS})$ and: $P(E_1, E_2, E_3) = P(E_1, E_2)$

Because both *MZ-twins* and *DZ-twins* share ($E_1$) and because both siblings and *DZ-twins* share the same genetic relationship with each other but only twins share the early environments, we expect that:

$$P(E_1 \mid MZ_{MS}) = P(E_1 \mid DZ_{MS}); \quad P(E_1 \mid S_{MS}) = P(E_1); \quad \text{and:} \; P(MS, E_2 \mid E_1, DZ_{MS}) = P(MS, E_2 \mid E_1, S_{MS})$$

In this case: $P(MS \mid DZ_{MS}) = P(MS, E_1, E_2 \mid DZ_{MS}) = P(E_1 \mid MZ_{MS}) * P(MS, E_2 \mid E_1, DZ_{MS})$

and: $P(MS \mid S_{MS}) = P(MS, E_1, E_2 \mid S_{MS}) = P(E_1) * P(MS, E_2 \mid E_1, DZ_{MS})$

so that: $\{P(MS \mid S_{MS}) / P(MS \mid DZ_{MS})\} * P(E_1 \mid MZ_{MS}) = P(E_1)$

And therefore: $P(MS \mid IG_{MS}) = \{P(MS \mid S_{MS}) / P(MS \mid DZ_{MS})\} * P(MS \mid MZ_{MS})$

#### 1.1. Penetrance Adjustments for different Population Partitions

***Conclusions:*** 1. The penetrance-adjustment for the shared early environment of *MZ*-twins is similar for both (*Male/Female*) and (*H+/H−*) partitions.

***Argument:*** Using the above relationship, from the Canadian population data set (*Table 2; Main Text*), we have already estimated the value of $\{P(MS \mid IG_{MS})\}$ as:

$$P(MS \mid IG_{MS}) = (2.9 / 5.4) * 0.25 = 0.25 / 1.86 = 0.134$$

In the more limited data from the HLA partition (*Table 2; Main Text*), this estimate becomes:

$$P(MS \mid IG_{MS}) = (2.9 / 5.4) * 0.30 = 0.30 / 1.86 = 0.161$$

Although these adjustments provide an estimate for the impact of the *IU* and early post-natal environment considering the population as a whole, the same adjustment may not be appropriate for every partition of that population. Using the data in *Table 2* (*Main Text*), however, certain partition-specific adjusted penetrance values can also be estimated and are quite similar for both partitions (*see below*).

#### 1.1a. Penetrance Adjustments for the (*H+/ H−*) Partition

***Conclusions:*** $P(MS \mid H+, IG_{MS}) = P(MS \mid H+, MZ_{MS}) / 1.84$

$P(MS \mid H-, IG_{MS}) = P(MS \mid H-, MZ_{MS}) / 1.87$

***Argument:*** Using the data in *Table 2* (*Main Text*), we can define two parameters $(a \geq 1)$ and $(b \geq 1)$ such that:

$$P(MS \mid H+, IG_{MS}) = P(MS \mid H+, MZ_{MS}) / a$$
$$P(MS \mid H-, IG_{MS}) = P(MS \mid H-, MZ_{MS}) / b$$

In this case we can deconstruct the term $\{P(MS, H+ \mid IG_{MS})\}$ in two different ways:

$$P(MS, H+ \mid IG_{MS}) = P(H+ \mid IG_{MS}) * P(MS \mid H+, IG_{MS}) = 0.43 * 0.31 / a = 0.133 / a$$

and: $$P(MS, H+ \mid IG_{MS}) = P(MS \mid IG_{MS}) * P(H+ \mid MS, IG_{MS}) = 0.161 * 0.45 = 0.072$$

Combining these two equations leads to: $a = 0.133 / 0.072 = 1.84$

Similarly: $$P(MS, H- \mid IG_{MS}) = P(H- \mid IG_{MS}) * P(MS \mid H-, IG_{MS}) = 0.57 * 0.29 / b = 0.165 / b$$

and: $$P(MS, H- \mid IG_{MS}) = P(MS \mid IG_{MS}) * P(H- \mid MS, IG_{MS}) = 0.161 * 0.55 = 0.089$$

leading to: $b = 0.165 / 0.089 = 1.87$

#### 1.1b. Penetrance adjustments for the (*F/ M*) partition

***Conclusions:*** $P(MS \mid F, IG_{MS}) = P(MS \mid F, MZ_{MS}) / 1.82$

$P(MS \mid M, IG_{MS}) = P(MS \mid M, MZ_{MS}) / 2.0$

***Argument:*** Similar to the analysis (*above*), using the data in Table 2 (*Main Text*), we can, again, define two parameters $(a \geq 1)$ and $(b \geq 1)$ such that:

$$P(MS \mid F, IG_{MS}) = P(MS \mid F, MZ_{MS}) / a$$
$$P(MS \mid M, IG_{MS}) = P(MS \mid M, MZ_{MS}) / b$$

Again, we can deconstruct the term $\{P(MS, F \mid IG_{MS})\}$ in two different ways:

$$P(MS, F \mid IG_{MS}) = P(F \mid IG_{MS}) * P(MS \mid F, IG_{MS}) = 0.66 * 0.34 / a = 0.224 / a$$

and: $$P(MS, F \mid IG_{MS}) = P(MS \mid IG_{MS}) * P(F \mid MS, IG_{MS}) = 0.134 * 0.92 = 0.123$$

Combining these two equations leads to: $a = 0.224 / 0.123 = 1.82$

Similarly: $P(MS, M \mid IG_{MS}) = P(M \mid IG_{MS}) * P(MS \mid M, IG_{MS}) = 0.34 * 0.07 / b = 0.024 / b$

and: $P(MS, M \mid IG_{MS}) = P(MS \mid IG_{MS}) * P(M \mid MS, IG_{MS}) = 0.134 * 0.08 = 0.012$

leading to: $b = 0.024 / 0.012 = 2.0$

**2. Enrichment of Genotypes – Partitioning (*G*) into Subsets (*G1*) & (*G2*)**

**2a. The 1st Enrichment**

***Conclusions:*** $P(MS \mid G1) = P(MS \mid G2)$ if and only if: $P(G1 \mid MS, G) = P(G1 \mid G)$

$P(MS \mid G1) > P(MS \mid G2)$ if and only if: $P(G1 \mid MS, G) > P(G1 \mid G)$

. ***Argument:*** The subset (*G*) can be partitioned into two mutually exclusive subsets (*G1*) and (*G2*), such that: $P(G) = P(G1) + P(G2)$ or: $1 = P(G1 \mid G) + P(G2 \mid G)$

For the 1st enrichment stage, we can define two constants, (*a*) and (*b*) such that:

$$P(MS \mid G1) = a * P(MS \mid G) \quad \text{and:} \quad P(MS \mid G2) = b * P(MS \mid G)$$

where (*a*) and (*b*) are related such that:

$$P(MS \mid G) = P(MS, G1 \mid G) + P(MS, G2 \mid G)$$

or: $1 = a * P(G1 \mid G) + b * P(G2 \mid G)$

and, thus: $a = 1$ if and only if: $b = 1$

also: $a > 1$ if and only if: $b < 1$ (and *vice versa*)

and finally: $P(MS \mid G1) / P(MS \mid G2) = (a / b)$

Moreover, for any partition:

$$P(G1 \mid MS, G) = P(G1, MS, G) / P(MS, G) = P(G1, G) * P(MS \mid G, G1) / P(MS, G)$$
$$= P(G1 \mid G) * a * P(MS \mid G) / P(MS \mid G) = a * P(G1 \mid G)$$

And similarly: $P(G2 \mid MS, G) = b * P(G2 \mid G)$

so that: $P(G1 \mid MS, G) / P(G2 \mid MS, G) = (a / b) * P(G1 \mid G) / P(G2 / G)$

Consequently, when: $a = b = 1$ then: $P(G1 \mid MS, G) = P(G1 \mid G)$ and: $P(G2 \mid MS, G) = P(G2 \mid G)$

However, if: $\sigma_X^2 > 0$ ; then there must be at least one partition, for which both (*G1*) and (*G2*) are non-empty and (*suitably defined*), for which: $(a > 1 > b)$, and therefore:

$$P(G1 \mid MS, G) > P(G1 \mid G) \quad \text{and:} \quad P(G2 \mid MS, G) < P(G2 \mid G)$$

Thus, any subset of more penetrant genotypes (*G1*) will be enriched in the $(MS, G)$ subset relative to a less penetrant subset (*G2*). Also, equally penetrant subsets will not be enriched relative to each other. Clearly, the reciprocal arguments also hold so that:

$$P(G1|MS,G) = P(G1|G) \quad \text{if and only if:} \quad P(MS|G1) = P(MS|G2)$$

and: $$P(G1|MS,G) > P(G1|G) \quad \text{if and only if:} \quad P(MS|G1) > P(MS|G2)$$

#### 2b. The 2nd Enrichment

***Conclusions:***

$$P(MS|G1,IG_{MS}) = P(MS|G2,IG_{MS}) \quad \text{if and only if:} \quad P(G1|MS,IG_{MS}) = P(G1|MS)$$

$$P(MS|G1,IG_{MS}) > P(MS|G2,IG_{MS}) \quad \text{if and only if:} \quad P(G1|MS,IG_{MS}) > P(G1|MS)$$

***Argument:*** Similar, for the 1st enrichment stage, we can define two constants, ($c$) and ($d$), such that: $$P(MS|G1,IG_{MS}) = c*P(MS|IG_{MS}) \quad \text{and:} \quad P(MS|G2,IG_{MS}) = d*P(MS|IG_{MS})$$

in which case: $$P(MS|IG_{MS}) = P(MS,G1|IG_{MS}) + P(MS,G2|IG_{MS})$$

or: $$1 = c*P(G1|MS) + d*P(G2|MS) = a*c*P(G1|G) + b*d*P(G2|G)$$

And, similar to the 1st enrichment stage, for the 2nd enrichment stage:

$$c = 1 \quad \text{if and only if:} \quad d = 1$$

and. also: $$c > 1 \quad \text{if and only if:} \quad d < 1$$

Moreover: $$P(G1|MS,IG_{MS}) = P(G1,MS,IG_{MS}) / P(MS,IG_{MS}) = c*P(G1|MS,G) = c*a*P(G1|G)$$

and: $$P(G2|MS,IG_{MS}) = d*P(G2|MS,G) = d*b*P(G2|G)$$

so that: $$P(G1|MS,IG_{MS}) / P(G2|MS,IG_{MS}) = (c/d)(a/b)\{P(G1)/P(G2)\}$$

#### 2c. Combining Enrichment Results

***Conclusions:*** Defining the additional parameters $(g, p, r, s, \ \& \ t)$ then:

1. $t = (a/b)\{p/(1-p)\}$
2. $x = p(x_1) + (1-p)(x_2)$
3. $(r/s)(a/b) = (c/d)$

***Argument:*** For simplicity of notation, in addition to those parameters $(a,b,c, \ \& \ d)$ defined in *#2a* and *#2b* (*above*) and $(x, x', x_1, x_1', x_2, \ \& \ x_2')$ already defined in *Proposition #1* (*Main Text*), we will define (*for this and the next two sections*) five additional parameters: $(g, p, \ r, \ s \ \& \ t)$ such that:

$$g = P(G); \quad p = P(G1|G); \quad r = (x_1')/(x_1); \quad s = (x_2')/(x_2); \quad \text{and:} \quad t = P(G1|MS)/P(G2|MS)$$

If the set $\{X\}$ – *see Proposition #1* (*Main Text*) – is bimodal based on the partition of ($G$) into high-penetrance ($G1$) and low-penetrance ($G2$) subsets, then from *#2a* (*above*):

$$P(G1|MS,G)/P(G2|MS,G) = (a/b)*P(G1|G)/P(G2/G)$$

or, equivalently: $t = (a/b)\{p/(1-p)\}$ (*Equation #1*)

and also: $P(MS \mid G) = P(MS)/g = x = p(x_1) + (1-p)(x_2)$

(*Equation #2*)

In addition, following the logic of *Proposition #1* (*Main Text*), if the set $\{X\}$ is bimodal as posited, then the penetrance values for the subsets ($G1$) and ($G2$), considered separtately, must be unimodal and each will conform to the upper quadratic solution such that:

$1 \le r < 2$ and, also: $1 \le s < 2$

Moreover: $(r/s) = \dfrac{(x_1')/(x_1)}{(x_2')/(x_2)} = (x_1'/x_2')(x_2/x_1) = (c/d)(b/a)$

so that: $(r/s)(a/b) = (c/d)$ (*Equation #3*)

where: $0.5 < (r/s) < 2$

**3. Proposition #1: Further Considerations**

**3a. Quadratic Considerations**

***Conclusions:*** 1. $x_1 = \dfrac{x + \sqrt{x^2 - \{1 + (r/s)(1-p)/p\}\{(x^2 - xx'(1-p)/s\}}}{p + (r/s)(1-p)}$

2. $x_2 = \dfrac{x - \sqrt{x^2 - \{1 + (s/r)p/(1-p)\}\{(x^2 - xx'p/r\}}}{(1-p) + (s/r)p}$

***Argument:*** Restating *Equation #2* (*see #2c above*):

$x = p(x_1) + (1-p)(x_2)$

so that: $x_2 = [x - p(x_1)]/(1-p)$ (*Equation #4a*)

In addition: $x' = P(MS \mid G, IG_{MS}) = P(MS, G1 \mid G, IG_{MS}) + P(MS, G2 \mid G, IG_{MS})$

therefore: $P(MS, G1 \mid G, IG_{MS}) = P(G1 \mid IG_{MS}) * (x_1') = P(G1 \mid MS) * (x_1')$

where: $P(G1 \mid MS) = P(G1, MS)/P(MS) = p(x_1)/x$

similarly: $P(G2 \mid MS) = (1-p)(x_2)/x$

so that: $xx' = pr(x_1)^2 + (1-p)s(x_2)^2$

or: $(x_2)^2 = [xx' - pr(x_1)^2]/(1-p)s$ (*Equation #4b*)

Consequently, we have two different estimates for $(x_2)^2$ – i.e., *Equations 4a and 4b, above*. Combining these two estimates yields:

$$[\{x - p(x_1)\}/(1-p)]^2 = (x_2)^2 = \{xx' - pr(x_1)^2\}/(1-p)s$$

or: $$[\{x - p(x_1)\}]^2 = (x_2)^2 = \{xx' - pr(x_1)^2\}(1-p)/s = \{xx'(1-p)/s\} - \{(r/s)p(1-p)(x_1)^2\}$$

and, finally: $$x^2 - 2xp(x_1) + p^2(x_1)^2 - xx'(1-p)/s + (r/s)p(1-p)(x_1)^2 = 0$$

Rearrangement, yields a quadratic equation in ($x_1$) such that:

$$\{p^2 + (r/s)p(1-p)\}(x_1)^2 - \{2xp\}(x_1) + \{x^2 - xx'(1-p)/s\} = 0$$

This, in turn, is solved (for $x_1$) by the Equation:

$$x_1 = \frac{x + \sqrt{x^2 - \{1 + (r/s)(1-p)/p\}\{(x^2 - xx'(1-p)/s\}}}{p + (r/s)(1-p)} \qquad (\textit{Equation\#5a})$$

In turn, solving for ($x_2$), yields:

$$x_2 = \frac{x - \sqrt{x^2 - \{1 + (s/r)p/(1-p)\}\{(x^2 - xx'p/r\}}}{(1-p) + (s/r)p} \qquad (\textit{Equation \#5b})$$

**3b. The Lower Solution**.

***Conclusions:*** $\forall\{P(MS|G) < x'/2\} : P(G1) \leq 0.05$

$\forall\{P(MS|G1) \leq 5 * P(MS|G2)\} : P(G) \leq 0.15$

$\forall\{P(G) = 1\} : P(MS|G1)/P(PMS|G2) > 40$

and: $\forall\{P(G) = 1\} : P(PMS|G2, IG_{MS}) < 0.0042$

***Argument:*** These quadratic solutions (*Equations 5a & 5b*) depend upon the values of four unknown variables *(e.g., r,s,p, & g)*. In addition, the values for $\{x', x_1', x_2'$ and: $P(MS)\}$ are based upon observation and, as such, subject to error. Therefore, these quadratic solutions cannot be solved uniquely. Nevertheless, a range of possible parameter values can be explored iteratively, using parameter combinations that cover (for each unknown parameter) their entire possible ranges, taking into account possible errors in our observations, and incorporating certain constraints on possible solutions. For example, for *Equation #5a* (*above*), the solution, ($x_1$), cannot be imaginary and the denominator cannot equal zero. Considering the partitions based on either gender or *HLA*-status, the plausible paramterer ranges for ($g$) and ($gp$) are:

$$0.0001 \leq g = P(G) \leq 1; \text{ and: } 0.0001 < gp = P(G1, G) < 0.5$$

For the Lower Solution to apply requires that: $0 < x_2 < x < x'/2$ – i.e., above this range the Upper Solution applies. Moreover, an acceptble solution cannot not be imaginary. Also for any solution, it must be that: $(x_1' > x')$ and that all estimated probabilities are within the range of zero to one. The observed value of $(x')$ was considered acceptible if it was within (±25%) of its observed value of: $x' = 0.134$. The observed value of $(x_2')$ was considered acceptible was less than or equal to twice is observed value in men (i.e., $x_2' \leq 0.0335 * 2 = 0.067$). The estimate for *P*(*MS*) was judged acceptible if it more than 85% and less than twice its observed value (i.e., $0.0026 \leq P(MS) \leq 0.006$). Because the Lower Solotion already indicates a bimodal distribution of the set {*X*}, the distribution for the penetrance values of the (*G*1) and and (*G*2) subsets must be unimodal (*see Methods*). As a result, the possilbe ranges for the (*r* and *s*) parameters are:

$$1 \leq r < 2; \quad \text{and:} \quad 1 \leq s < 2$$

Considering parameter-value combinations, which span the entirity of these ranges, we find that :

$$\forall \{P(MS \mid G) < x'/2\} : P(G1) \leq 0.05$$

$$\forall \{P(MS \mid G1) \leq 5 * P(MS \mid G2)\} : \ P(G) \leq 0.15$$

$$\forall \{P(G) = 1\} : \ P(MS \mid G1) / P(PMS \mid G2) > 40$$

and: $$\forall \{P(G) = 1\} : \ P(MS \mid G2, IG_{MS}) < 0.0042$$

Thus, although Lower Solutions exist for which, $\{P(G) = 1\}$, none of these simultaneously match the constraints placed by the observed the values of $(x', x_1', \ \& \ x_2')$ in the partitions based on gender or *HLA*-status. We conclude, therefore, that the circumstance of $\{P(G) = 1\}$ is not possible. Moreover, as suggested by our selection of the above ranges, any acceptable solution must also conform to the circumstances both for the susceptible persons, generally, and also for those of men and women considered separately (*see #4 below*). Finally, in earlier iterations of this analysis [3,4,49,50], we defined the subset-(*G*) differently – i.e., $\forall G_i \in G : \ P(MS \mid G_i) \geq P(MS)$. We note that, in the present analysis, using our new defnitions, our older definition corresponds to defining only members of the (*G*1)-subset as being "genetically susceptible" to MS.

### 4. Genetic Susceptibility including Women and Men – Variance Considerations

***Conclusions:***

1. The set {*X*} has a bimodal distribution
2. $0.163 \leq P(MS \mid F, G) \leq 0.187$
3. $0.029 \leq P(MS \mid M, G) \leq 0.034$
4. $0.23 \leq P(F \mid G) \leq 0.28$

5. $4.8 \leq P(MS \mid F,G) / P(MS \mid F,G) \leq 6.4$

6. $0.080 \leq P(G) < 0.094$

***Argument:*** Following the notation, definitions, and logic of *#2 & #3 (above)* and of *Proposition #1* (*see Main Text*), we can define a partition of the ($G$) subset into susceptible women $\{(G1) = (F,G)\}$ and susceptible men $\{(G2) = (M,G)\}$.

The set $\{X\}$ of penetrance values for members of the ($G$)-subset (*Proposition #1* ; *Main Text*) is clearly bimodal. Thus, from the observational data in *Table #2* (*Main Text*):

$$x_1' = 0.34 >> 0.067 = x_2' \; ; \quad X^2 = 8.5 \; ; \; p = 0.0035$$

Because we are only considering the possibility of either unimodal or bimodal distributions for the set $\{X\}$, the sets ($G1$) and ($G2$), considered separately, therefore will be unimodal and, thus, meet conditions for the Upper Solution. Naturally, the fact that the bimodal nature of the set $\{X\}$ is revealed by the gender partition does not indicate that biologic criteria for determining the bimodality is necessarily gender. Rather, it only means that the gender partition must substantially capture this criteria. For example, using an entirely hypothetical circumstance, suppose that the relevant biological criteria was a blood estrogen level above a certain critical value. In such circumstance, only rare women may fail to meet this criteria and, conversely, only rare men may meet it. In this case, absent knowing the actual criteria, the bimodality will appear to be based on gender. Nevertheless, in such a circumstance, partitioning ($G$) based on the actual criteria will only serve to increase the separation between the expected penetrance of the ($G1$) and ($G2$) subsets.

Therefore, using logic identical to that of *Proposition #1* for the ($M / F$) partition, using the estimated adjustments for the similar early environment of twins for these two subgroups (*see #2 above*), and using the data provided in *Table 2* (*Main Text*), for the Upper Solution, it follows that:

$$0.093 < x_1 \leq 0.187 \qquad (\textit{Equation \#6a})$$

and: $$0.017 < x_2 \leq 0.034 \qquad (\textit{Equation \#6b})$$

The proportion of MS patients who are women from *Table 2* (*Main Text*) is 66%. For the WTCCC data this number is 72%. From the study of Orton and colleagues [54] out of Canada, in the most recent epoch, the percentage of MS patients who are women is 76%. Therefore, using the smallest (i.e., the most conservative) of these estimated gender imbalances, using the above ranges for men and women, and from the definition of the ($G$)-subset, we can estimate that:

$$P(MS,G \mid F) = P(MS \mid F) = \{P(F \mid MS) * P(MS)\} / P(F) \approx \{0.66 * 0.003\} / 0.5 = 0.004$$

Because: $P(G \mid F) = P(MS,G \mid F) / P(MS \mid F,G)$

Therefore: $0.021 = 0.004 / 0.187 \leq P(G \mid F) < 0.004 / 0.093 = 0.043$

And similarly: $0.060 = 0.002 / 0.034 \leq P(G \mid M) < 0.002 / 0.017 = 0.118$

These possible ranges for men and women don't overlap and therefore, at a minimum, the upper limit for the proportion of women in the ($G$)-subset (predicted by the upper solution) can be estimated.

Thus: $P(G \mid F) / P(G \mid M) = P(F \mid G) / P(M \mid G) = p / (1-p) < 0.043 / 0.060 = 0.717$

where: $1 + P(F \mid G) / P(M \mid G) = 1 / P(M \mid G) = 1 / (1-p) < 1 + 0.717 = 1.717$

so that: $p = P(F \mid G) < 0.717 / 1.717 = 0.42$

However, in undertaking this calculation, there are four serious concerns. First, in using the above limits for $(x_1$ and: $x_2)$, we are positing and an extreme and tri-modal distribution for the set {$X$} – i.e., not the unimodal or bimodal distributions under consideration in this manuscript. Thus, this calculation, envisions a distribution, in which half of the women have a penetrance of slightly greater than zero and the other half have a uniform penetrance of $(x_1')$ – i.e. women have the maximum variance possible – and in which each of the men has exactly the same penetrance of $(x_2')$, which is intermediate between these two extremes – i.e. men have a zero variance.

Second, such an extreme distribution seems very unlikely, especially in the circumstance where partitioning the ($G$)-subset by a different MS-associated characteristic – i.e., *HLA*-status (*see #5, below*) – doesn't even give a hint of the bimodal nature of {$X$}.

Third, it is not possible that the variance of penetrance values for the $(F,G)$-subset to be at its maximum value. Thus, because $(x' < x_1')$, then the maximum variance for the ($G1$)-subset $- (x_1'/2)^2$ – exceeds the maximum total variance possible for the entire ($G$)-subset – $(x'/2)^2$. Consequently, the lower limit for the value of ($x_1$) in *Equation #6a* – i.e., at its maximum possible variance – must be too low.

And fourth, some of the maximum possible variance in the {$X$} set must be accounted for just by the separation of $(x_1)$ from $(x_2)$. To see this, suppose that the {$X$} set is bimodal as described (*above*) for gender. And further suppose that, of the ($m$) members of the ($G$) subset, $(ii = 1,2,3,...,g1)$ belong to the high-penetrance ($G1$)-subset and $(jj = 1,2,3,...,g2)$ belong the mutually exclusive low-penetrance ($G2$)-subset. In this case, $(p = g1/m)$ and the variance of the set {$X$} – as described in *Proposition #1* (*Main Text*) – is:

$$\sigma_X^2 = E(x_i - x)^2 = (1/m)\sum_{i=1}^{m}(x_i - x)^2 = (1/m)\sum_{ii=1}^{g1}(x_{ii} - x)^2 + (1/m)\sum_{jj=1}^{g2}(x_{jj} - x)^2$$

For the set ($G1$), following a standard development of variance relationships [55], this becomes:

$$(1/m)\sum_{ii=1}^{g1}(x_{ii}-x)^2 = (1/m)\sum_{ii=1}^{g1}[(x_{ii}-x_1)+(x_1-x)]^2 = (1/m)\sum_{ii=1}^{g1}(x_{ii}-x_1)^2 + (1/m)\sum_{ii=1}^{g1}(x_1-x)^2$$

or: $$(1/m)\sum_{i=1}^{g1}(x_{ii}-x)^2 = (g1/m)\sigma_{x1}^2 + (g1/m)(x_1-x)^2 = p\sigma_{x1}^2 + p(x_1-x)^2$$

Similarly: $$(1/m)\sum_{jj=1}^{g2}(x_{jj}-x)^2 = (1-p)\sigma_{x2}^2 + (1-p)(x_2-x)^2$$

Consequently: $$\sigma_X^2 = \{p\sigma_{x1}^2 + (1-p)\sigma_{x2}^2\} + \{p(x_1-x)^2 + (1-p)(x_2-x)^2\}$$ *Equation #4*

Thus, part of the set {$X$} variance is accounted for by the term: $\{p(x_1-x)^2 + (1-p)(x_2-x)^2\}$. This will also cause the lower limit for the value of ($x_1$) to be higher than expressed in that *Equation #6a*. For example, we can define the residual variance ($R$) as:

$$R = \sigma_X^2 - p(x_1-x)^2 - (1-p)(x_2-x)^2$$

where: $\sigma_X^2 = x(x'-x)$ ; $\sigma_{x1}^2 = x_1(x_1'-x_1)$ ; $x = px_1 + (1-p)x_2$

As before, the upper limit for $P(G1|G) = P(F|G)$ can be established using the maximum value for $(x_2)$ – i.e., where: $\sigma_{x2}^2 = 0$ – and the minimum value for $(x_1)$ – i.e., where $(\sigma_{x1}^2)$ accounts for all of the residual varaince ($R$). Under these circumstances, together with this definitions and these relationships, we have two equations in only two unknown (i.e., unobserved) parameters $(p$ and: $x_1)$, which can be solved uniquely. Thus, using the relationship defined previously that:

$$t = P(G1|MS)/P(G2|MS)$$

and using the upper-solution (*Proposition #1, Main Text*), at the boundry, $(x_1$ and: $p)$ become:

$$p = t(x_1/x_2)/\{1+t(x_1/x_2)\}$$ *Equation #3; re-expressed*

and: $$x_1 = \frac{(x_1') + \sqrt{(x_1')^2 - 4R}}{2}$$ where: $\sigma_{x1}^2 \le R$

In turn, these two equations can be solved iteratively by inserting our initial lower-limit estimate of $(x_1 = 0.093)$ into the 1st equation and then estimating a new lower limit $(x_1)$ from the 2nd equation, which takes into account the condition that: $(\sigma_{x1}^2 \le R)$. This new lower-limit estimate can then be inserted into the 1st equation an the process repeated until the two estimates for $(x_1)$ converge and are identical.

Initially assuming that the entire residual variance (*R*) is confined to penetrance values in the $(G1)$-subset (i.e., $\sigma^2_{x2} = 0)$, and using *Equation #5*, this process converges on the solution that:

$$x_1 > 0.145 \ ; \quad p < 0.31 \ ; \text{ and: } (a/b) > 4.3$$

Permitting the variance $(\sigma^2_{x2})$ to increase to its maximum possible value of: $\{x_2(x_2' - x_2)\}$ only alters these conclusions with respect to $\{p$ and: $(a/b)\}$ such that:

$$x_1 > 0.145 \ ; \quad p < 0.18 \ ; \text{ and: } (a/b) > 8.7$$

Nevertheless, both of these solutions, once again, describe the circumstances of extreme trimodal (or multimodal) distributions. Therefore, to avoid this, the distribution of penetrance values in (*G*1) and *G*2) must each meet the requirements for a unimodal distribution (*see Proposition #1, Main Text*) and, in this circumstance, the solution to these two equations becomes:

$$0.163 \le x_1 \le 0.187 \ ; \ 0.029 \le x_2 \le 0.034 \ ; \ 0.23 \le p \le 0.28 \ ; \text{ and: } 4.8 \le (a/b) \le 6.4$$

Also, from this we conclude that:

$$0.011 = 0.002/0.187 \le P(G,F) < 0.002/0.163 = 0.013$$

and: $$0.030 = 0.001/0.034 \le P(G,M) < 0.001/0.029 = 0.034$$

so that: $$0.041 \le P(G) = P(G,F) + P(G,M) < 0.047$$

The estimate from *Table 2* (*Main Text*) for the quantity $\{P(MS \mid M, IG_{MS})\}$, is based on only two observations, and, thus, this estimate seems likely to be the least reliable of any in the *Table.* However, even if this estimated penetrance were doubled, there would still be an excess of men in the (*G*)-subset such that:

$$0.37 \le P(F \mid G) \le 0.44$$

and also: $$0.026 \le P(G) \le 0.030$$

**5. Genetic Susceptibility for the (*H*+) / (*H*−) Partition**

***Conclusion:*** $P(G \mid H+) \approx 3.35 * P(G \mid H-)$

***Argument:*** From the WTCCC it is apparent that there is considerable enrichment of (*H*+) status during the 1st enrichment stage when moving from the general population to an MS population.

Thus: $$P(MS,G \mid H+) / P(MS,G \mid H-) = 3.35$$

This relationship can be expressed alternatively as:

$$\{P(G \mid H+) / P(G \mid H-)\} * \{P(MS \mid G,H+) / P(MS \mid G,H-)\} = 3.35$$

What this re-expression makes it clear that the enrichment of ($H$+)-status in MS can occur in one, or both, of two possible ways. First, ($H$+) membership could make membership in the ($G$)-subset more likely than it is for the ($H$−) subset – i.e., it is due to an impact on the ratio of: $P(G \mid H+) / P(G \mid H-)$. Second, members of the $(G, H+)$ subset may have a greater penetrance for MS than members of the $(G, H-)$ subset – i.e., it is due to an impact on the ratio of: $P(MS \mid G, H+) / P(MS \mid G, H-)$.

As discussed in *#4* (*above*), the set $\{X\}$, based on the partition of the ($G$)-subset by gender, is clearly bimodal. Therefore, there is a continuing enrichment of women during both the 1st and the 2nd stage of enrichment as demonstrated (*see Table 2; Main Text*) by the following relationships:

$$P(F \mid MS) / P(F) = 1.32 \approx 1.39 = P(F \mid MS, IG_{MS}) / P(F \mid MS)$$

By contrast, the enrichment of ($H$+)-status disappears during the (observable) 2nd stage such that:

$$P(H+ \mid MS) / P(H+) = 1.79 > 1.05 = P(H+ \mid MS, IG_{MS}) / P(H+ \mid MS)$$

However, if either: $P(F, H+ \mid G) > P(F, H- \mid G)$ or: $P(F, H- \mid G) > P(F, H+ \mid G)$

then the group with the greater proportion of women will be enriched (due to the considerably greater penetrance in women – *see #4, above*) during the 2nd enrichment stage. This should be true even if ($H$+) has little impact by itself. The enrichment should be even greater if ($H$+) has an additional impact although, in fact, there is little observational evidence for this. Thus, based on the limited data of *Table 2* (*Main Text*):

$$P(F, H+ \mid MS) = 0.34 \approx 0.35 = P(F, H- \mid MS)$$

and: $$P(MS \mid F, H+, IG_{MS}) = 0.21 \approx 0.24 = P(MS \mid F, H-, IG_{MS})$$

Taken together, these two observations suggest that there is little, or no, enrichment of ($H$+) status during the 2nd stage in women. Consequently, gender-status must be approximately balanced within the two *HLA*-subgroups such that:

$$P(F, H+, G) \approx P(F, H-, G)$$

In which case: $P(G \mid F, H+) * P(F, H+) \approx P(G \mid F, H-) * P(F, H-)$

Using the WTCCC control data, this relationship translates to:

$$P(G \mid F, H+) * 0.23 \approx P(G \mid F, H-) * 0.77$$

or: $$P(G \mid F, H+) \approx 3.35 * P(G \mid F, H-)$$

Similarly, in WTCCC controls: $P(H+|F) \approx P(H+|M)$, so that for men also:

$$P(G|M,H+) \approx 3.35 * P(G|M,H-)$$

Therefore: $P(G|H+) \approx 3.35 * P(G|H-)$

Consequently, the large majority of the enrichment of ($H+$) status in MS seems to be due to the fact that ($H+$)-membership makes membership in the ($G$)-subset more likely than it is for ($H-$) membership. By contrast, ($H+$)-status seems to have very little impact on penetrance.

### 6. Environmental Considerations in MS-Pathogenesis for Men and Women

#### 6a. Re-expressing Penetrance at *Time-period #1* in terms of *Time-period #2*

***Conclusions:*** 1. $Zw_1 = P(MS,E|G,F)_1 = \{P(F|MS)_1 / P(F|MS)_2\} * C * (Zw_2)$

2. $Zm_1 = P(MS,E|G,M)_1 = \{P(M|MS)_1 / P(M|MS)_2\} * C * (Zm_2)$

***Argument:*** From the definitions (*Main Text*) of ($Zm_1$), ($Zm_2$) and of the subsets ($G$) and ($E$):

$$P(MS,E,G,F) = P(MS,F)$$

and, thus: $Zw_2 = P(MS,E|G,F)_2 = P(MS,F)_2 / P(G,F) = P(MS)_2 * P(F|MS)_2 / P(G,F)$

Because, by the definition of ($C$) – see *Main Text*: $P(MS)_1 = C * P(MS)_2$

Therefore: $Zw_1 = P(MS,E|G,F)_1 = C * P(MS)_2 * P(F|MS)_1 / P(G,F)$

And, thus: $Zw_1 = (Zw_2) * C * \{P(F|MS)_1 / P(F|MS)_2)\}$

Similarly: $Zm_1 = (Zm_2) * C * \{P(M|MS)_1 / P(M|MS)_2)\}$

#### 6b. Determining the Limiting Values for Penetrance

***Conclusions:*** 1. $d = (Zw_2) * \{1 - [P(F|MS)_1 / P(F|MS)_2] * C * e^{-1}\} / (1 - e^{-1})$

2. $c = (Zm_2) * \{1 - [P(M|MS)_1 / P(M|MS)_2] * C * e^{-1}\} / (1 - e^{-1})$

***Argument:*** From the *Main Text*, using the scale ($a^{app} = R*a$) for *women*:

$$Zw_2 = P(MS,E|G,F)_2 = d * \{1 - e^{-(a1^{app} + 1 - \lambda w)}\}$$
$$Zw_1 = P(MS,E|G,F)_1 = d * \{1 - e^{-(a1^{app} - \lambda w)}\}$$

These two equations can be re-arranged to yield:

$$(Zw_2 - d) / d = -\{e^{-(a1^{app} - \lambda w)}\} * e^{-1}$$
$$(Zw_1 - d) / d = -\{e^{-(a1^{app} - \lambda w)}\}$$

Dividing these two equations yields: $(Zw_2 - d) / (Zw_1 - d) = e^{-1}$

and with re-arrangement, this equation yields: $d = P(MS \mid G,E,F) = \{Zw_2 - Zw_1 * e^{-1}\} / (1 - e^{-1})$

Substituting into this last equation for ($Zw_1$) from (*#5a*; *above*) yields:

$$d = (Zw_2) * \{1 - [P(F \mid MS)_1 / P(F \mid MS)_2] * C * e^{-1}\} / (1 - e^{-1})$$

Analogously: $$c = P(MS \mid G,E,M) = \{Zm_2 - Zm_1 * e^{-1}\} / (1 - e^{-1})$$

and: $$c = (Zm_2) * \{1 - [P(M \mid MS)_1 / P(M \mid MS)_2] * C * e^{-1}\} / (1 - e^{-1})$$

**6c. Assessing the Environmental Threshold for MS in Men and Women**

***Conclusions:*** 1. $\lambda = \ln[(c/d)(Zw_2 - d)/(Zm_2 - c)]$

2. $\forall C > 0 : \lambda > 0$

***Argument:*** Because, the scales for the response-curves for women and men are proportional, the response curve for women − along the *x-axis* in (*a*) units − will be stretched or compressed, depending upon the value of (*R*), compared to the response-curve when: $R = 1$ (*Fig. 3; Main Text*). The threshold (*x-intercept*) occurs at $\{(a, Zm) = (\lambda m, 0)\}$ for men and $\{(a, Zw) = (\lambda w, 0)\}$ for women. By the definitions of (*E*) and (*a*), one of these thresholds must occur at $\{(a, Z) = (0,0)\}$. However, these thresholds need not be the same so we define the difference in threshold between women and men as: $(\lambda = \lambda w - \lambda m)$ so that, if women have a higher threshold than men: $(\lambda > 0)$. On the ($a^{app}$) scale, the *x-intercept* always occurs at $\{a^{app} = \lambda w = 0\}$, and, thus, $(\lambda w)$ is independent of *R*. The condition $(R = 1)$, therefore, can be used to estimate $(\lambda w)$, and responses at the second time-point can be expressed as:

$$Zw_2 = P(MS,E \mid G,F)_2 = d * \{1 - e^{-(a1+1-\lambda w)}\} = d * \{1 - e^{-(a1+1-\lambda-\lambda m)}\} \qquad (women)$$
$$Zm_2 = P(MS,E \mid G,M)_2 = c * \{1 - e^{-(a1+1-\lambda m)}\} \qquad (men)$$

With re-arrangement, these equations *yield*:

$$(Zw_2 - d)/d = -e^{-(a1+1-\lambda m)}\} * e^{\lambda}$$
$$(Zm_2 - c)/c = -e^{-(a1+1-\lambda m)}\}$$

Dividing these two equations yields: $(c/d) * [Zw_2 - d)/(Zm_2 - c)] = e^{\lambda}$

So that: $\lambda = \ln\{(c/d) * [Zw_2 - d)/(Zm_2 - c)]\}$

Using the, from the values of $(Zw_2 = 0.181)$ and $(Zm_2 = 0.031)$ provided in the *Main Text*, and defining two expressions in ($C$) – ($K_M$) and $K_F$) – such that:

$$K_M = [P(M \mid MS)_1 / P(M \mid MS)_2] * C * e^{-1} = 0.483 * C$$

and: $$K_F = [P(F \mid MS)_1 / P(F \mid MS)_2] * C * e^{-1} = 0.332 * C$$

Then the equations for ($c$) and ($d$) in *#6b* (*above*) can be re-expressed as:

$$c = Zm_2 * (1 - K_M) / (1 - e^{-1})$$

and: $$d = Zw_2 * (1 - K_F) / (1 - e^{-1})$$

Using the *above* equation for $(\lambda)$, applying *#6d* (*below*), and assuming that, in Canada: $C > 0.50$, then

$$\forall C > 0.50:\ 0.37 < \lambda < 4.67\ ;\ \text{and, in fact:}\ \ \forall C > 0 : \lambda > 0$$

**6d. Assessing the increase in MS-prevalence for Canada (1945 – 1980)**

***Conclusions:*** 1. $C = P(MS)_1 / P(MS)_2 < 0.76$

2. $P(MS)_2 > (1/C) * P(MS)_1 > 1.32 * P(MS)_1$

***Argument:*** From *#6a* and *#6b* (*above*):

$$Zm_1 = (Zm_2) * C * P(M \mid MS)_1 / P(M \mid MS)_2\}$$

and: $$c = P(MS \mid G, E, M) = \{Zm_2 - Zm_1 * e^{-1}\} / (1 - e^{-1})$$

Substituting the 1st equation into the 2nd, and noting that $\{Zm_2 < c\}$, yields:

$$Zm_2 < \{Zm_2 - [P(M \mid MS)_1 / P(M \mid MS)_2] * C * (Zm_2) * e^{-1}\} / (1 - e^{-1})$$

Dividing through by ($Zm_2$) and with re-arrangement this yields:

$$C < \{1/(1 - e^{-1}) - 1\} / \{[P(M \mid MS)_1 / P(M \mid MS)_2] * e^{-1} / (1 - e^{-1})\} = 0.76$$

Therefore: $$P(MS)_2 = (1/C) * P(MS)_1 = 1.32 * P(MS)_1$$

**6e. Proportional vs. Non-proportional Hazard-rates for Men and Women**

***Conclusions:*** 1. The limiting-values of (***c***) and (***d***) are the same in either case

2. Non-proportional hazards require separate graphs for men and women

3. Non-proportional hazards imply different environmental factors

4. Proportional hazards imply similar environmental factors

***Argument:*** From the separate definitions for the response curves in woman and men (*described above in #6b*), it follows directly that the limiting-values (***c***) and (***d***) depend only upon the independent variables ($a$) and ($a^{app}$) respectively. However, non-proportionality suggests that there is no known relationship between these two variables and, therefore, that no further comparative information can be gleaned. The response curves can be graphed separately but, because the relationship between the scales of the two is not known, they can't be placed on the same graph. Nevertheless, if the relationship between these two scales could be defined in some other manner, it might be possible to develop comparative information. In addition, non-proportionality would suggest that the environmental factors, which contribute to $P(E)$, are different for men and women.

By contrast, the assumption of proportional hazard rates directly implies that women have a higher environmental threshold than men *(Fig. 3; see #6c above*) and such a circumstance suggests that men and women are responding to the same environmental events. Otherwise, if the necessary environmental factors were different for men and women, there must be some specific environmental conditions that favor MS-development in women over men and, in such a case, there could be no consistent difference in threshold. Rather, the existence of a threshold difference between men and women suggests that any gender-specific differences in MS-development depend only upon the degree (not the kind) of exposure. For example, perhaps, men become susceptible with a lesser degree of vitamin D deficiency or with EBV infection occurring over a broader age-range compared to women.

**7. Uniqueness of Susceptible Genotypes in the Population**

An early GWAS of the WTCCC data identified a set of 102 non-*MHC* SNPs, for which one of the two SNP-variants at each location was significantly (and reliably) associated with MS [13]. Thus, including the "risk" haplotype ($H+$) in the *HLA* Class II region of the *MHC*, 103 "risk" locations were identified [13]. Among control subjects, there were, on average, 31 of these locations at which subjects were homozygous for the "non-risk" *SNP* variant, 32 locations at which subjects were homozygous for the "risk" *SNP*-variant, and 40 locations at which control subjects were heterozygous. By contrast, among cases, these numbers were 29, 34, and 40 (respectively). Even if one considers individuals who are either homozygous or heterozygous for the "risk" *SNP*-variant at all locations to have the same genotype, there are still a huge number of possible combinations. Thus, in this circumstance there are:

$$\left(\frac{103}{74}\right) = 3.4 * 10^{25}$$

possible combinations. Nevertheless, it is clear, first, that heterozygotes and homozygotes carry different "risks" for some locations (e.g., $H+$), second, that many of the associated loci have multiple alleles (e.g., the *MHC*), and finally, that more than 200 genetic loci are now known to be MS-associated [5-14,24]. Each of these facts will hugely increase the number of possible genetic combinations of the MS-associated loci.

With fewer than {$10^{10}$} people in the entire world, it seems almost certain that everyone (except monozygotic twins) will have a unique genotype when considering this entire collection of susceptibility locations. Indeed, we used the 30,248 individuals in the WTCCC dataset to test this hypothesis [13]. The first 102 WTCCC-identified non-*MHC* SNPs [13] were ordered by the strength of their MS-association (i.e., by the magnitude of their respective *ORs*). Considering heterozygotes and homozygotes to be separate genotypes and considering only 20 of the strongest MS-associated haplotypes – together with the (*H*+)-genotype in the Class II region – no genotype (including both cases or controls) had more than 2 representations in the WTCCC and considering just 85 of the 103 regions (including *H*+), everyone had a unique genotype. Similarly, considering heterozygotes and homozygotes to be the same genotype and considering only 40 of the 103 regions, no genotype had more than two representations in the WTCCC and considering just 86 regions, everyone had a unique genotype. Clearly, neither including in the analysis the more than 200 loci now-identified MS-associated *SNP*s [14], nor analyzing, at these susceptibility loci, specific MS-associated *SNP*-haplotypes rather than single *SNP*s [23,24], will alter this conclusion. Everyone in the WTCCC has a unique genotype considering all of their MS-associated loci.

We also analyzed the WTCCC data for every possible combination of three "risk" haplotypes at these 102 locations, together with the state of the (*H*+)-genotype, with regard to their MS-association. There were 960 combinations (homozygous, heterozygous, or either), together with a heterozygous state at the (*H*+) locus, that had an *OR*, which significantly exceeded that found by considering the (*H*+) locus by itself. By contrast, there were 7009 such combinations, together with a homozygous state at the (*H*+) locus, that had an *OR*, which significantly exceeded that when considering the (*H*+) locus by itself. Nevertheless, in both circumstances, there was little consistency. Counting the number of cases having each 4-locus combination for heterozygous (*H*+) individuals, yielded: (mean = 51; range: 14 – 582) and, for homozygous (*H*+) individuals, it yielded: (mean = 112; range: 22 – 338). Also, these estimates steadily decreased with each additional locus included in the combination. These observations become more striking when one considers just the 100 most significant MS-associated combinations for individuals heterozygous or homozygous for (*H*+) separately. In the heterozygous group, the number of cases having each combination is: (mean = 34; range: 16 – 92), whereas the number of cases not having each combination is far greater: (mean = 4,272; range: 4,684 – 4,760). Similarly, in the homozygous group, the number of cases having each combination is: (mean = 54; range: 31 – 130), whereas the number of cases not having each combination is, again, far greater: (mean = 753; range: 677 – 776). Thus, it seems clear that, although certain combinations increase the likelihood of (*G*)-subset membership, the actual combinations that do this are quite heterogeneous, and only a small proportion of genetically susceptible individuals (who actually develop MS) share even the same 4-locus genetic combination. This finding indicates that genetic susceptibility to MS is largely idiosyncratic.
